# SLC7A5 is required for citrulline-dependent growth in arginine limited conditions

**DOI:** 10.1101/2024.06.27.600869

**Authors:** Kyle N. Dunlap, Austin Bender, Alexis Bowles, Alex J. Bott, Jared Rutter, Gregory S. Ducker

## Abstract

Tumor cells must optimize metabolite acquisition between synthesis and uptake from their surroundings. The tumor microenvironment is characterized by hypoxia, lactate accumulation, and depletion of many circulating metabolites, including amino acids such as arginine. We performed a metabolism-focused functional screen using CRISPR/Cas9 in a melanoma cell line to identify pathways and factors that enable tumor growth in an arginine-depleted environment. Our screen identified the SLC-family transporter SLC7A5 as required for growth, and we hypothesized that this protein functions as a high-affinity citrulline transporter. Citrulline, an essential precursor to arginine synthesis, is present in human serum at 40 μM and supports localized arginine synthesis across diverse tissues. Using isotopic tracing experiments, we show that citrulline uptake and metabolism are dependent upon expression of this transporter. Pharmacological inhibition of SLC7A5 blocks growth in low arginine conditions across a diverse group of cancer cell lines. Loss of SLC7A5 reduces tumor growth and citrulline import in a mouse tumor model. Overall, we identify a conditionally essential role for SLC7A5 in arginine metabolism as a mediator of citrulline uptake, and we propose that SLC7A5-targeting therapeutic strategies in cancer may be especially effective in the context of arginine limitation.

**Key Points:**

- SLC7A5 is required for proliferation in arginine-free conditions when citrulline is present.
- SLC7A5 loss impairs arginine metabolism.
- Citrulline import is uniquely dependent on SLC7A5.
- Small molecule inhibitors of SLC7A5 can be paired with senolytic drugs to drive apoptosis.
- *SLC7A5* knockout decreases citrulline import in a xenograft model.

## Introduction

The distribution of nutrients via circulation enables individual cells and tissues to either acquire or synthesize nutrients depending upon their niche and function. Diet provides a steady stream of all proteinogenic amino acids, but humans remain capable of synthesizing approximately half of them, giving cells multiple strategies for acquiring amino acids^1^. Despite this theoretical flexibility, non-essential amino acids are limiting in proliferative contexts such as immune cell expansion^2,3^ or tumor growth^4^. Strategies to boost or inhibit amino acid acquisition through transport or synthesis have been elucidated for multiple tumor and immune cell interactions^5,6^. Arginine is both a proteinogenic amino acid and a precursor for biochemical processes such as polyamine synthesis and nitric oxide production and signaling. Blocking arginine availability has major effects on tumor growth and T-cell function in multiple systems^7–11^. It has been shown that T-cells may be able to circumvent the loss of environmental arginine through the uptake of citrulline from the serum to fuel the de novo synthesis of arginine^12^.

Citrulline is a neutral non-proteinogenic amino acid whose primary physiological function is to serve as an essential intermediate in the urea cycle to detoxify ammonia in the liver^13^. Citrulline and aspartate are combined into arginosuccinate by the enzyme Argininosuccinate Synthase 1 (ASS1). Fumarate is then cleaved from arginosuccinate to generate arginine by the action of Argininosuccinate Lyase (ASL). Loss of citrulline catabolism by mutations in *ASS1* blocks urea cycle function, leading to citrullinemia type 1, a buildup of citrulline and ammonia in the blood^14^. ASS1 and ASL are widely expressed, and citrulline metabolism enables arginine synthesis in peripheral tissues^15^. The total arginine synthesis capacity is large as citrulline is sufficient to sustain serum arginine and ornithine levels, even in the presence of exogenously administered arginases^16^.

*ASS1* is both up and down-regulated within tumors,^17^ suggesting that cancers face differing pressures on arginine availability and placing citrulline acquisition as a key node in tumor metabolism. *ASS1* expression is repressed by epigenetic silencing in multiple cancer types, including melanoma^18^, mesothelioma^19^, and hepatocellular carcinoma^7,20^. Loss of ASS1 can aid tumor growth by sparing aspartate for nucleotide synthesis^7,21^. Unsurprisingly, these tumors are exquisitely sensitive to arginine depletion, leading to clinical trials seeking to exploit this metabolic vulnerability by using an arginine deiminase enzyme therapy (ADI-PEG20) to catabolize circulating arginine into citrulline and the ammonium ion^22^. A recent phase II clinical trial in mesothelioma showed increased overall survival in patients treated with ADI-PEG20, although *ASS1* expression was not a patient-specific variable in this study^23^. Re-expression of *ASS1* could enable tumor cells to escape arginine-depletion treatment by utilizing citrulline^24^. More broadly, overexpression of *ASS1* may be a signal that the tumor microenvironment is depleted of arginine and that citrulline acquisition and utilization may represent a targetable metabolic vulnerability.

In this study, we sought to characterize systemic citrulline metabolism and quantify citrulline consumption across tissues. To understand the necessary metabolic networks required for citrulline metabolism, we performed a functional genomics screen using CRISPR. This screen identified *SLC7A5*, which encodes a neutral amino acid transporter, as essential for citrulline to rescue growth when arginine is limited. We demonstrate that this essentiality is due to the loss of citrulline import and de novo arginine synthesis, as well as the specificity of citrulline for SLC7A5 compared to other annotated amino acid targets. We show that a small molecule inhibitor of SLC7A5 can pair with arginine starvation in *ASS1*-high cells to inhibit cell growth. Finally, we show in vivo that SLC7A5 deficient tumors have reduced growth and attenuated citrulline import.

## Results

### A functional genomics screen identifies SLC7A5 as required for growth on citrulline

Citrulline is a human plasma metabolite used for arginine synthesis (Fig. 1A), but its role in metabolism outside of the liver and the urea cycle has been a subject of ongoing investigation. To measure citrulline concentration and flux in circulation, we infused [1-^13^C]-citrulline and [^13^C_6_]-arginine over three hours into fasted 9-week-old mice through a surgically implanted jugular vein catheter. We measured the fasting serum concentration of citrulline to be 66 ± 5 μM, and arginine to be 103 ± 15 μM, consistent with prior reports from both mice and humans^25^ (Fig. 1B). Upon infusion (parameters listed in methods), both tracers reached steady state enrichments (28% isotope enrichment for citrulline and 14% for arginine) during the infusion period (Fig. S1A). From serum isotopic enrichment at steady state, we quantified the endogenous rate of appearance (*R_a_*) of citrulline and arginine. Our quantified arginine production flux (12.5 ± 1.7 nmol/g/min) was more than four times that of citrulline (2.6 ± 0.3 nmol/g/min) (Fig. 1C). As circulating concentrations of arginine and citrulline were constant during this infusion period, we can also assume that total tissue disposal fluxes (*R_d_*) were equivalent to *R_a_* and higher for arginine than citrulline. [1-^13^C]-citrulline infusion led to the labeling of serum arginine, indicating active arginine synthesis from citrulline (Fig. S1B). [^13^C_6_]-arginine infusion resulted in M+5 labelled citrulline that linearly increased during the infusion, indicating systemic nitrous oxide synthetase activity^26,27^ (Fig S1C). These data show that in mice, systemic arginine and citrulline metabolism are closely interconnected.

**Figure 1:**
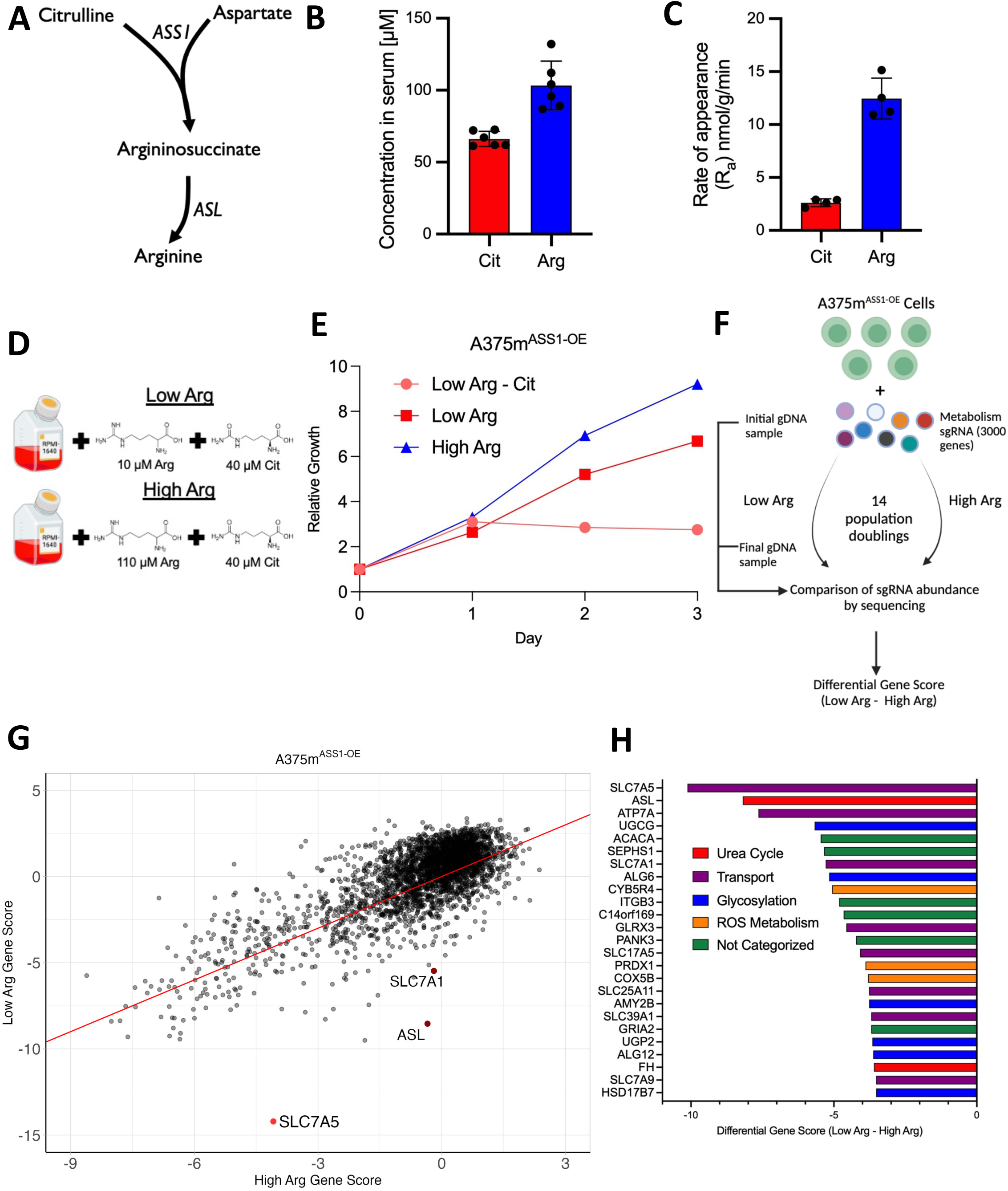
A functional genomics screen identifies SLC7A5 as required for growth on citrulline. A: Schematic of citrulline synthesis into arginine. B: Concentrations of arginine and citrulline in mouse serum. (mean ± SD, n = 5). C: Rates of appearance of arginine and citrulline as determined from a steady state labeling fraction after a 3-hour infusion. (mean ± SD, n = 5). D: Media conditions used for the CRISPR screen. E: Growth of A375m^ASS1-OE^ cells in Low Arg, High Arg, and Low Arg - Cit media. Data is expressed in terms of relative growth, where the readings at day 0 are normalized to 1. (mean ± SD, n = 4). F: Schematic of the CRISPR-based screen to identify metabolic genes required for growth in low arginine. G: Gene scores in cells grown in Low Arg vs. High Arg. SLC7A5, the top hit, is highlighted in red. SLC7A1 (arginine transporter) and ASL (arginosuccinate lyase) are also highlighted. The red line is the equation y = x passing though (0,0) to highlight differential essentiality. H: Top 25 genes scoring as selectively essential in Low Arg vs. High Arg. Genes linked to glycosylation are in blue, the urea cycle in red, transport in purple, Reactive Oxygen Species (ROS) metabolism in orange, and other genes in green.

To identify cellular mechanisms regulating arginine synthesis from citrulline, we designed a functional genomics screen to identify genes involved in citrulline uptake and metabolism. We generated a cell line that was dependent upon citrulline metabolism for growth in arginine-depleted conditions by overexpressing ASS1 in the naturally *ASS1*-low melanoma cell line A375m (A375m^ASS1-OE^) (Fig. S1D). To establish reliance upon de novo arginine synthesis, we cultured these cells in media with limiting levels of arginine (Fig. 1D). We supplemented RPMI media formulated without arginine with 40 µM citrulline, the concentration of citrulline contained in Human Plasma Like Media (HPLM)^25^. We then added back either 10 µM Arg (Low Arg), a near-starvation concentration, or 110 µM Arg (High Arg), the concentration of Arg in HPLM and similar to what we observed in mouse serum^28^ (Fig. 1B). We performed growth assays and observed a slight reduction in growth in the A375m^ASS1-OE^-Low Arg condition compared to the A375m^ASS1-OE^-High Arg condition, but without citrulline present, growth was fully suppressed after one doubling, consistent with the exhaustion of pre-existing arginine and citrulline (Fig. 1E). To test whether ASS1 activity could fully support cell growth, we cultured cells in supraphysiological levels of citrulline (200 µM) and rescued proliferation back to High Arg levels (Fig. S1E). To confirm that citrulline metabolism was increased by ASS1 overexpression, we quantified citrulline and arginine consumption from media using A375m^ASS1-OE^ and ASS1 knockout (A375m^ASS1-KO^) cell lines. A375m^ASS1-OE^ cells consumed more citrulline and less arginine in High Arg conditions compared to A375m^ASS1-KO^ cells (Fig. S1F). ASS1 overexpression enabled citrulline consumption in Low Arg media compared to the A375m^ASS1-KO^ cells. Furthermore, ornithine, another non-proteinogenic amino acid member of the urea cycle, could not rescue growth in arginine-limiting conditions, demonstrating a lack of ornithine transcarbamylase (OTC) enzyme expression in our cell line (Fig. S1G). This result supports prior reports that OTC is only expressed in the liver and parts of the intestine and not expressed in the vast majority of cultured cell lines (Fig. S1H, A375 cell line highlighted in magenta)^29,30^.

Having established the arginine synthesis phenotype in our engineered cell lines, we then performed a functional genomics screen using CRISPR/Cas9 in culture media with differing defined arginine and citrulline concentrations. We employed a previously published metabolism focused library that consists of ∼24,000 sgRNAs targeting ∼3,000 genes and enzymes relating to human metabolism and small molecule transporters (8 sgRNA/gene) and 50 control sgRNAs in a Cas9-expressing lentiviral vector^31,32^. We transduced A375m^ASS1-OE^ cells with the sgRNA library, passaged the pool of cells in either Low Arg or High Arg media for 14 population doublings, and quantified sgRNA abundances before and after the experiment (Fig. 1F). We utilized MAGeCK software^33^ to calculate gene scores based on the essentiality of the gene for cellular fitness. As expected, most genes scored similarly between the Low and High Arg conditions (Fig. 1G). Consistent with our system design, genes within the urea cycle scored as highly essential in Low Arg conditions. The second most differentially essential gene was *ASL*, which encodes the enzyme arginosuccinate lyase, the final step in generating arginine from citrulline (Fig. 1G, 1H, S1I). A canonical arginine transporter, *SLC7A1*^7^, was also a top hit (Fig. 1G, 1H, S1I). However, the most differentially essential gene between the Low Arg and High Arg conditions was *SLC7A5* (Fig. 1G, 1H, S1I, S1J). It ranked as the most essential gene in the A375m^ASS1-OE^-Low Arg condition, and the #182 most essential gene in the A375m^ASS1-OE^-High Arg condition (Fig. S1I). SLC7A5 (also referred to as LAT1) is a sodium- and pH-independent amino acid antiporter^34^. It is a well-annotated high-affinity transporter of large, neutral amino acids such as leucine, phenylalanine, and tryptophan^34–36^. As citrulline is neutral at physiologic pH and the most differentially essential gene in our screen, we hypothesized that SLC7A5 was responsible for its transport in low arginine conditions.

### SLC7A5 is required for citrulline uptake, metabolism, and growth in arginine-free media

To validate the essential role of SLC7A5 for cell growth in low arginine conditions, we generated two clonal *SLC7A5* knockout lines in our A375m^ASS1-OE^ cell background using CRISPR/Cas9 (Fig. 2A). A375m^ASS1-OE^ WT and *SLC7A5* KO cells were grown over 4 days in arginine-free RPMI supplemented with equimolar amounts of either arginine or citrulline (110 µM each). A modest but significant reduction in doubling time was observed in some *SLC7A5* knockout clones compared to A375m^ASS1-OE^ WT cells when cultured with arginine (Fig 2B, S2C). However, when cultured on citrulline in place of arginine, *SLC7A5* KO cells were unable to proliferate at all (Fig. 2B). Re-expression of *SLC7A5* cDNA rescued growth in media containing citrulline (Fig. S2A, S2B). To examine if this phenotype was broadly applicable across cell types, we used CRISPR-Cas9 to knock out *SLC7A5* in the naturally ASS1-high (Fig. S5C) MCF7 breast cancer cell line (Fig 2C)^30^. Like A375m^ASS1-OE^ cells, knockout of SLC7A5 inhibited growth of MCF7 cells when they were cultured with citrulline in place of arginine (Fig. 2D).

**Figure 2:**
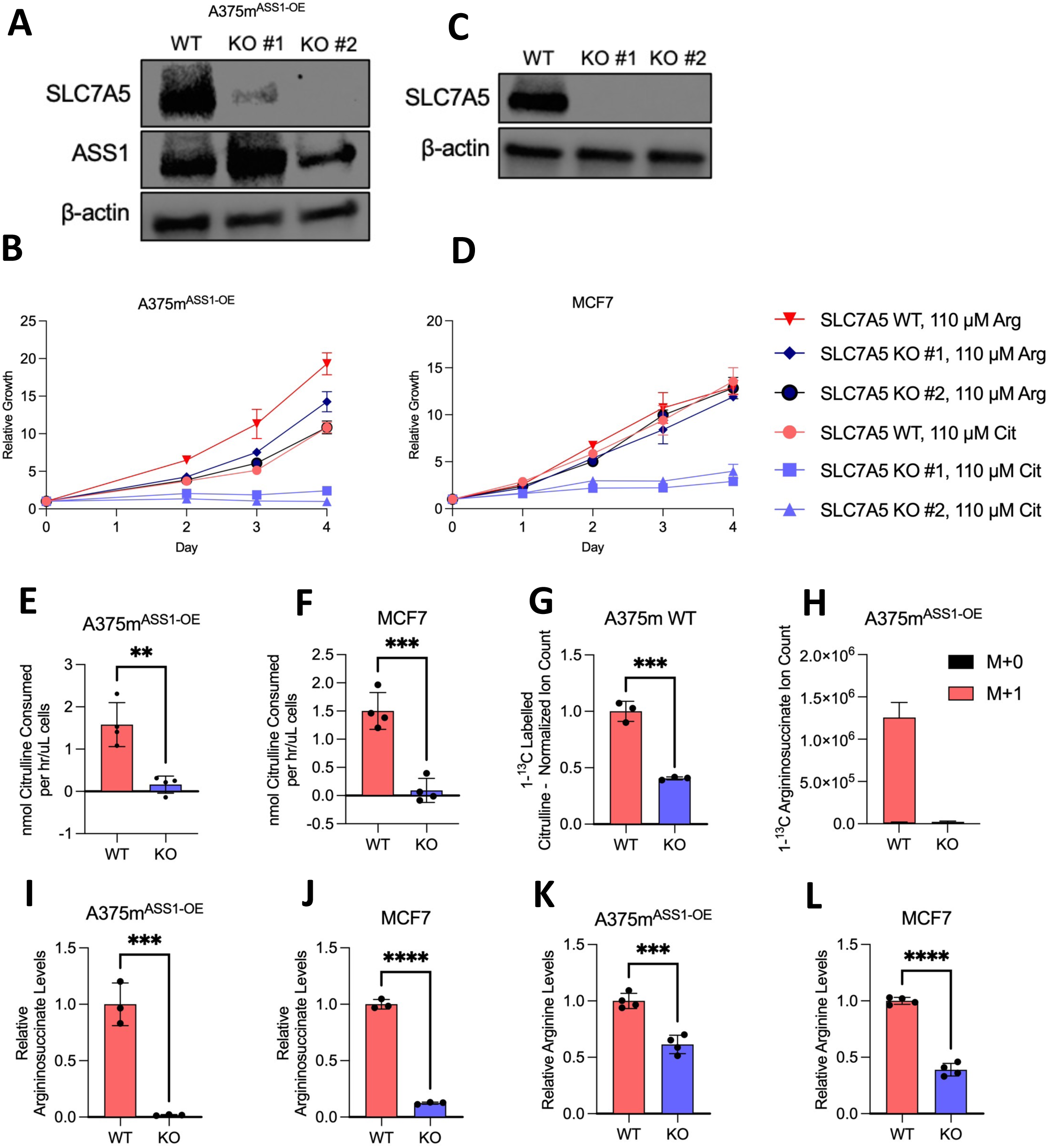
SLC7A5 is required for citrulline uptake, metabolism, and growth in arginine-free media. A: Immunoblot of two clones of the SLC7A5 KO and ASS1-OE in the A375m^ASS1-OE^ (melanoma) cell line. B: A375m^ASS1-OE^ and SLC7A5 KO cells were grown in indicated media. The base media was RPMI without arginine supplemented with 110 μM arginine or citrulline was added as indicated. Media was refreshed on day 2. (mean ± SD, n = 4). C: Immunoblot of two clones of the SLC7A5 KO and ASS1-OE in the MCF7 (breast cancer) cell line. D: MCF7 SLC7A5 WT and KO cells were grown in indicated media. Media was refreshed on day 2. (mean ± SD, n = 4). E: Citrulline consumption from the media with over 48-hour period, A375m^ASS1-OE^ and SLC7A5 KO. Media: RPMI + 110 μM Arg + 110 μM Cit. (mean ± SD, n = 4). ** p < .01, unpaired, two-tailed t-test. F: Citrulline consumption from the media with SLC7A5 WT and KO over a 48-hour period, MCF7 WT and SLC7A5 KO. Media: RPMI + 110 μM Arg + 110 μM Cit. (mean ± SD, n = 4). *** p < .001, unpaired, two-tailed t-test. G: Uptake of [1-^13^C]-Citrulline over a 15-minute period in A375m WT and SLC7A5 KO cells. Media: RPMI + 110 μM Arg + 40 μM Cit. (mean ± SD, n = 3). *** p < .001, unpaired, two-tailed t-test. H: Ion count of unlabeled (M+0) and labeled (M+1) argininosuccinate. Cells were plated in RPMI + 110 μM [1-^13^C]-Citrulline for 16 hours and then metabolites were extracted and analyzed using LC-MS. I: Intracellular argininosuccinate levels in A375m^ASS1-OE^ cells after 24 hours of culture in RPMI + 110 μM Cit. (mean ± SD, n = 4). *** p < .001, unpaired, two-tailed t-test. J: Intracellular argininosuccinate levels in MCF7 cells after 24 hours of culture in RPMI + 110 μM Cit. (mean ± SD, n = 4). **** p < .0001, unpaired, two-tailed t-test. K: Intracellular arginine levels in A375m^ASS1-OE^ cells after 24 hours of culture in RPMI + 110 μM Cit. (mean ± SD, n = 4). *** p < .001, unpaired, two-tailed t-test. L: Intracellular arginine levels in MCF7 cells after 24 hours of culture in RPMI + 110 μM Cit. (mean ± SD, n = 4). **** p < .0001, unpaired, two-tailed t-test.

To understand whether our citrulline-dependent growth phenotype was related to the transport function of SLC7A5, we examined the metabolic consequences of SLC7A5 KO (KO #2 was used for all subsequent experiments) using liquid chromatography-mass spectrometry (LC-MS). SLC7A5 is a neutral amino acid antiporter that utilizes glutamine^37^, and intracellular glutamine levels were increased in SLC7A5 KO cell lines in arginine-free media (Fig. S2D, S2E). We next quantified the effects of SLC7A5 deletion on the total consumption of citrulline in arginine-free media. Citrulline uptake rate was calculated by measuring the media citrulline concentration over time and normalizing it to cell growth. Knockout of SLC7A5 fully blocked consumption of citrulline in both cell lines (Fig. 2E, 2F). Re-expression of *SLC7A5* cDNA rescued citrulline consumption (Fig. S2F). To compliment these measurements, we quantified cellular uptake of [1-^13^C]-citrulline in A375m WT (not overexpressing ASS1) cells over a period of 15 minutes and found that SLC7A5 KO cells show reduced uptake of labeled citrulline (Fig. 2G). We traced the ASS1-catalyzed conversion of citrulline to arginosuccinate using [1-^13^C]-citrulline. In A375m^ASS1-OE^ cells given [1-^13^C]-citrulline, we observed ^13^C isotope labeling of arginosuccinate that was no longer detectable in A375m^ASS1-OE^ SLC7A5 KO cells (Fig 2H). Total intracellular levels of argininosuccinate were also significantly reduced in SLC7A5 KO cells cultured in citrulline (Fig. 2I, 2J). Restoration of SLC7A5 rescued arginosuccinate levels (Fig S2G). Because of this defect in arginine synthesis, total intracellular arginine levels were significantly reduced in KO cells in both cell lines (Fig 2K, 2L). Arginine levels were increased when SLC7A5 was re-expressed (Fig. S2H). Collectively, these data suggest that SLC7A5 is essential for the uptake of citrulline and synthesis of arginine in cultured cell lines.

### Under physiological amino acid concentrations, citrulline uptake is uniquely dependent upon SLC7A5

SLC7A5 is a high-affinity transporter for neutral amino acids such as leucine and phenylalanine and has been reported to affect mTOR signaling due to its leucine transport activity^38,39^. To determine whether the growth defects we observed in low arginine conditions were confounded due to loss of transport of these amino acids, we performed growth assays with variable concentrations of these metabolites. We reasoned that if SLC7A5 was important for the uptake of these amino acids, A375m^ASS1-OE^ SLC7A5 KO cells would exhibit deficient growth when cultured in low levels of these amino acids. At physiological levels of leucine and phenylalanine, we failed to observe a growth defect in SLC7A5 KO cells (Fig. 3A, B; see dashed line). For leucine, we did not observe a genotype specific growth defect until levels were ∼1/8 (25 μM) of physiologic concentrations. For phenylalanine, the effect was seen at ∼1/3 of physiologic levels (25 μM). As we predicted based on published data, cells cultured in very low levels of leucine displayed both an increase in SLC7A5 expression and ATF4 expression confirming the high affinity transport role of SLC7A5 (Fig. S3A). In contrast, when we grew cells in media without arginine supplemented with citrulline over a range of concentrations both above and below physiological levels, we observed a large growth defect when SLC7A5 was absent (Fig 3C). Based on regression analysis, the amino acid concentration that reduced cell growth to half of maximal was less than 7 μM for leucine and phenylalanine in both WT and KO cells, while for citrulline, 220 μM was required to support half maximal growth in KO cell lines (Fig. S3B). Citrulline only rescued growth in SLC7A5 KO at ∼10 times the physiological concentration of this metabolite, implying the existence of a second, very low-affinity mechanism of citrulline transport. We confirmed these amino acid growth dependencies in a second cell line, MCF7 (Fig. 3D-F, S3C). These data suggest that SLC7A5 is essential for citrulline import at physiological concentrations, while other amino acid transporters can compensate for the transport of leucine and phenylalanine at serum concentrations of these amino acids.

**Figure 3:**
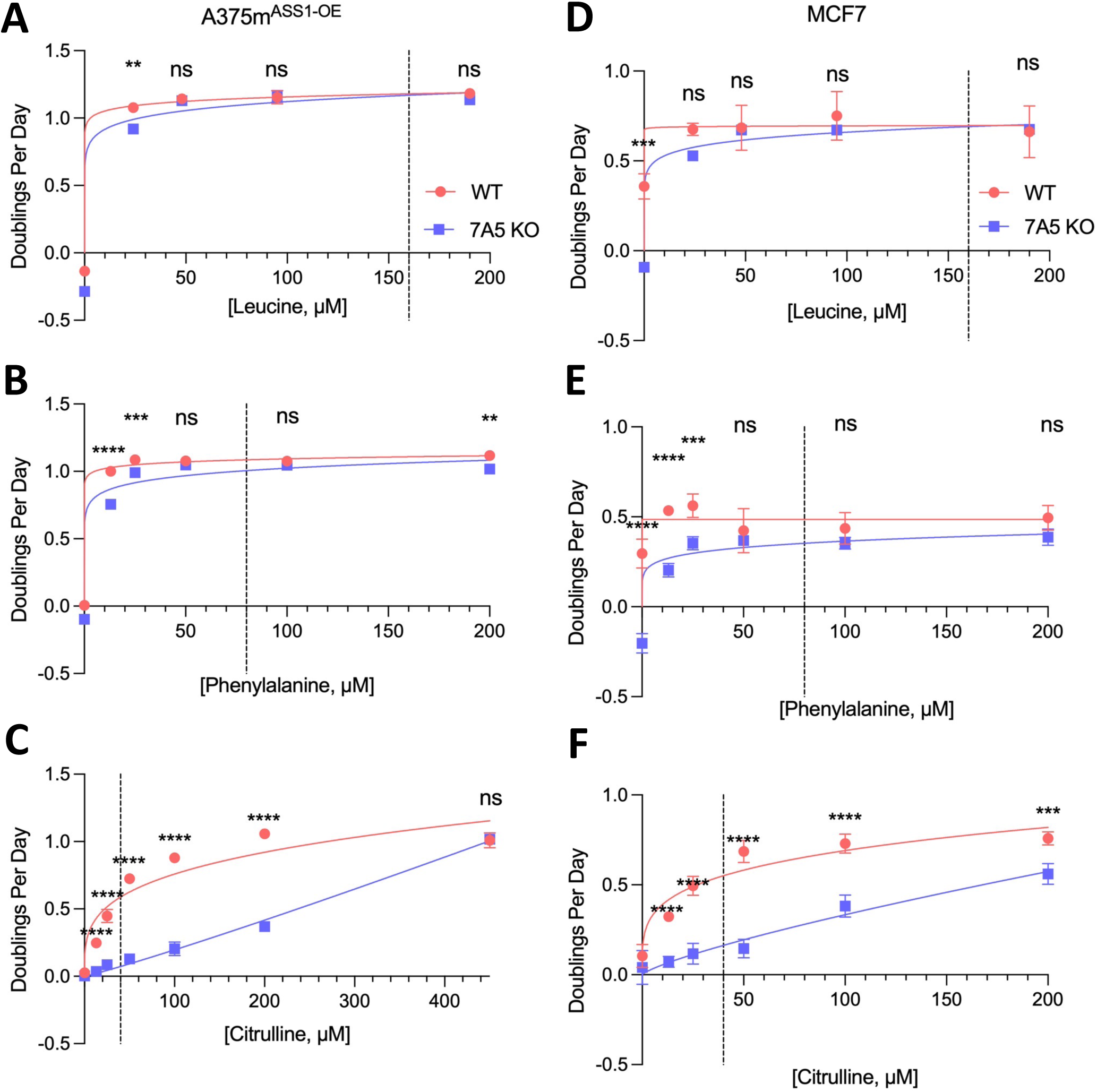
Under physiological amino acid concentrations, citrulline uptake is uniquely dependent upon SLC7A5. A: Growth of A375m^ASS1-OE^ WT and SLC7A5 KO cells in RPMI + variable amounts of leucine. The dashed line is 160 μM, the concentration of leucine in HPLM. Doublings per day data calculated after 3 days of growth in the respective media. Media was refreshed on day 2. (mean ± SD, n = 4). ** p < .01, unpaired, two-tailed t-test. The line of best fit for all graphs in this figure was calculated as Y = 10^(slope*log(X) + Yintercept). B: Growth of A375m^ASS1-OE^ WT and SLC7A5 KO cells in RPMI + variable amounts of phenylalanine. The dashed line is 80 μM, the concentration of phenylalanine in HPLM. Doublings per day data calculated after 3 days of growth in the respective media. Media was refreshed on day 2. (mean ± SD, n = 4). n.s. not significant, ** p < .01, *** p < .001, **** p < .0001, unpaired, two-tailed t-test. C: Growth of A375m^ASS1-OE^ WT and SLC7A5 KO cells in RPMI + variable amounts of citrulline. This media did not contain arginine. The dashed line is 40 μM, the concentration of citrulline in HPLM. Doublings per day data calculated after 3 days of growth in the respective media. Media was refreshed on day 2. (mean ± SD, n = 4). n.s. not significant, **** p < .0001, unpaired, two-tailed t-test. D: Growth curves of MCF7 WT and SLC7A5 KO cells in RPMI + variable amounts of leucine. The dashed line is 160 μM, the concentration of leucine in HPLM. Doublings per day data calculated after 4 days of growth in the respective media. Media was refreshed on day 2. (mean ± SD, n = 4). n.s. not significant, *** p < .001, unpaired, two-tailed t-test. E: Growth curves of MCF7 WT and SLC7A5 KO cells in RPMI + variable amounts of phenylalanine. The dashed line is 80 μM, the concentration of phenylalanine in HPLM. Doublings per day data calculated after 4 days of growth in the respective media. Media was refreshed on day 2. (mean ± SD, n = 4). n.s. not significant, *** p < .001, **** p < .0001, unpaired, two-tailed t-test. F: Growth curves of MCF7 WT and SLC7A5 KO cells in RPMI + variable amounts of citrulline. This media did not contain arginine. The dashed line is 40 μM, the concentration of citrulline in HPLM. Doublings per day data calculated after 4 days of growth in the respective media. Media was refreshed on day 2. (mean ± SD, n = 4). n.s. not significant, *** p < .001, **** p < .0001, unpaired, two-tailed t-test.

### *SLC7A5* and *ASS1* are upregulated in response to arginine starvation

If SLC7A5 was necessary for growth in arginine limited conditions, we asked if restricting arginine would lead to increased SLC7A5 expression. Arginine starvation in A375m (ASS1 WT) cells led to arginine stress as evidenced by increased transcript levels of ATF4, a transcription factor involved in the integrated stress response responsive to arginine^40^, and commensurate upregulation of transcripts of the arginine metabolic enzyme *ASS1* and the arginine transporter *SLC7A1* (Fig. 4A). *SLC7A5* transcripts were also significantly upregulated in response to this condition (Fig. 4A). In multiple *ASS1*-low cell lines, arginine starvation increased expression of both ASS1 and SLC7A5 protein (Fig. 4B). We next asked if a similar mechanism accounted for these increases. In A375m WT cells, we observed that loss of arginine in the presence of citrulline led to global increase in intracellular levels of amino acids, including neutral essential amino acids. (Fig. 4C, S4G, red vs blue bars). This result phenocopies an earlier study focusing on glutamine deprivation, where it was found that glutamine-starved environments have increased intracellular amino acid levels.^41^. This led us to ask whether *SLC7A5* and *ASS1* were regulated downstream of the general control nonderepressible 2 (GCN2) response to uncharged tRNA. Treatment with GCN2iB, a GCN2 inhibitor, largely normalized cellular amino acid levels (Fig. 4C, S4G, red vs pink bars). Inhibition of GCN2 blocked ATF4 expression in response to arginine starvation but did not impact SLC7A5 expression (Fig 4D). In response to arginine starvation, SLC7A5 is upregulated alongside known arginine metabolism genes, albeit through an ATF4-independent mechanism.

**Figure 4:**
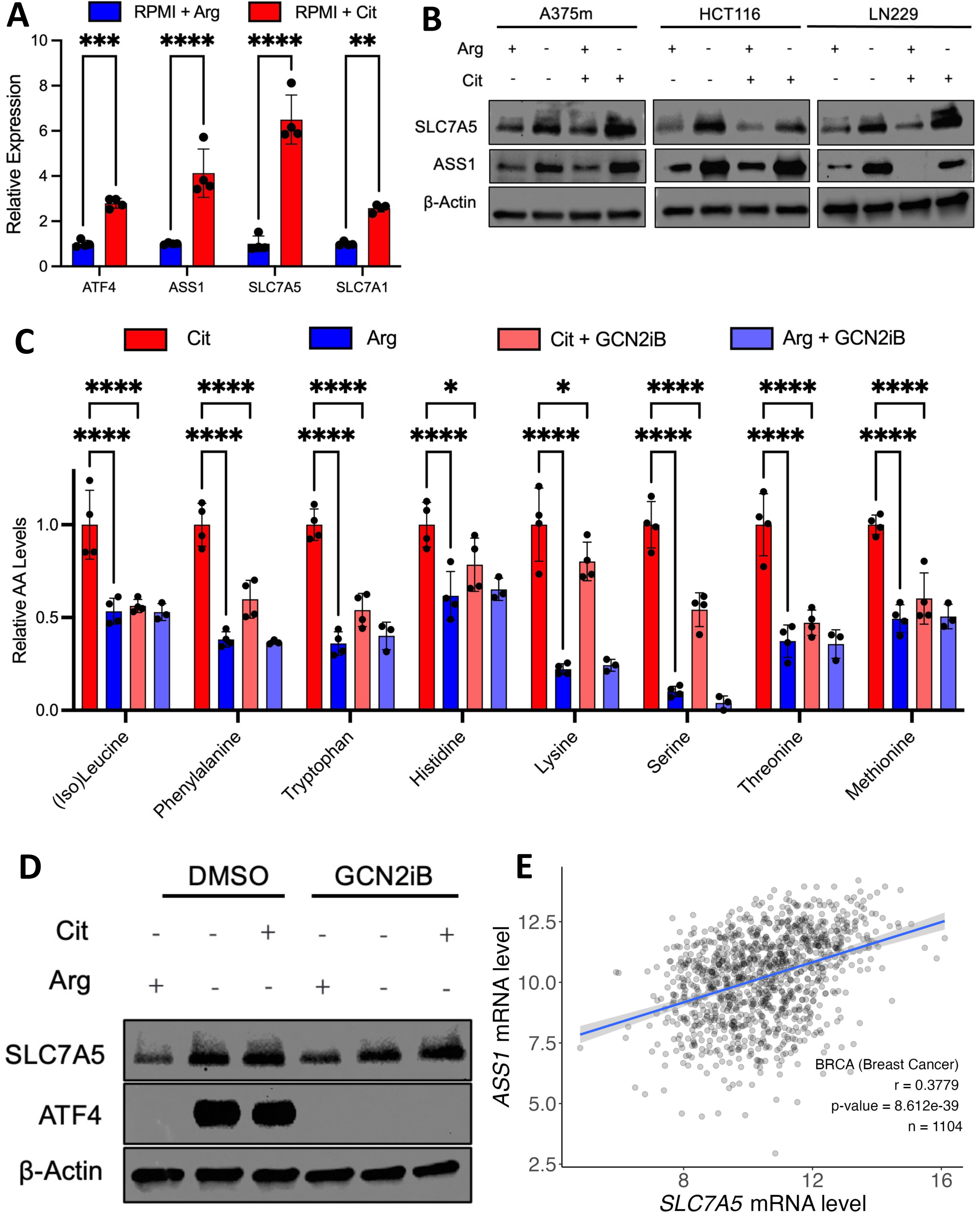
*SLC7A5* and *ASS1* are upregulated in response to arginine starvation. A: Relative transcript levels of *ATF4, ASS1, SLC7A5* and *SLC7A1* in A375m WT cells cultured in the indicated media as determined by qRT-PCR. *RPS2* was used as a normalizing gene. (mean ± SD, n = 4). ** p < .01, *** p < .001, **** p < .0001, 2-way ANOVA and multiple comparisons were corrected for using the Sidak method. B: Immunoblots of SLC7A5 and ASS1 from A375m, HCT116 and LN229 WT cells plated for 48 hours in indicated media conditions. C: Relative amino acid levels of cells plated for 24 hours in either RPMI + 110 μM Cit (Cit, red), RPMI + 110 μM Arg (Arg, blue), RPMI + 110 μM Cit + 1 μM GCN2iB (Cit + GCN2iB, salmon) or RPMI + 110 μM Arg + 1 μM GCN2iB (Arg + GCN2iB, periwinkle). (mean ± SD, n = 3-4). * = p < .05, **** = p < .0001, multiple comparisons-corrected 2-way ANOVA. D: Immunoblot of SLC7A5 and ATF4 in A375m cells plated for 48 hours in indicated media conditions in media containing indicated combinations of RPMI + 110 μM Cit or RPMI + 110 μM Arg. GCN2iB was dosed at 1 μM. E: Correlation between transcript levels of *SLC7A5* versus *ASS1* in breast cancer (BRCA) tumor transcript data taken from TCGA. The gray area around the linear regression line indicates the 95% confidence interval.

While SLC7A5 and ASS1 are both necessary for growth on citrulline, we asked how they cooperated to control citrulline metabolic fluxes. We assayed citrulline uptake across a panel of 6 cancer cell lines and in A375m^ASS1-OE^ cells. Citrulline uptake varied widely in media containing both citrulline and arginine among the cell lines and was closely associated with growth rate in media containing citrulline, but not in media containing arginine (Fig S4A, S4B). We then immunoblotted for ASS1 and SLC7A5 levels in our cell lines and observed a strong correlation between growth rate on citrulline and ASS1 but not SLC7A5 expression (Fig S4B, S4C). To explore whether SLC7A5 expression was necessary for arginine synthesis in human cancers, we examined the correlation between *SLC7A5* and *ASS1* mRNA expression. Data from cancerous breast tissue cataloged in The Cancer Genome Atlas (TCGA) shows significant co-expression between *SLC7A5* and *ASS1* among all breast cancers (Fig. 4E), of which strongest correlation occurred in the more aggressive Luminal B subtype (Fig. S4D), while Luminal A tumors showed a much weaker correlation (Fig. S4E). In kidney chromophobe cancer, a rare subtype that originates from intercalated cells of distal tubules^42^, a strong correlation was also observed (Fig. S4F).

### A small molecule inhibitor of SLC7A5 sensitizes cells to arginine deprivation

JPH203 is a potent and selective inhibitor of SLC7A5 currently in clinical trials for biliary tract cancer^43^. We asked whether treatment with JPH203 would selectively target cancer cells in arginine depleted conditions. JPH203 (1 μM) inhibited growth of six different cancer cell lines when arginine was restricted and citrulline provided (Fig. 5A, red dots). Importantly, at this dose, JPH203 did not affect growth of cells grown in arginine, indicating that this may be a citrulline-dependent phenotype and not a general result of blocking proteinogenic amino acid uptake (Fig. 5A, blue dots). To confirm that this effect is transport-specific, we added either a supraphysiological amount of citrulline (1 mM) or physiologic arginine (110 μM) to the media and found that they both rescue growth (Fig. 5B). As we observed in SLC7A5 KO cells (Fig 2), JPH203 treatment significantly reduced arginosuccinate and arginine levels in cells (Fig. 5C, 5D, S5A).

**Figure 5:**
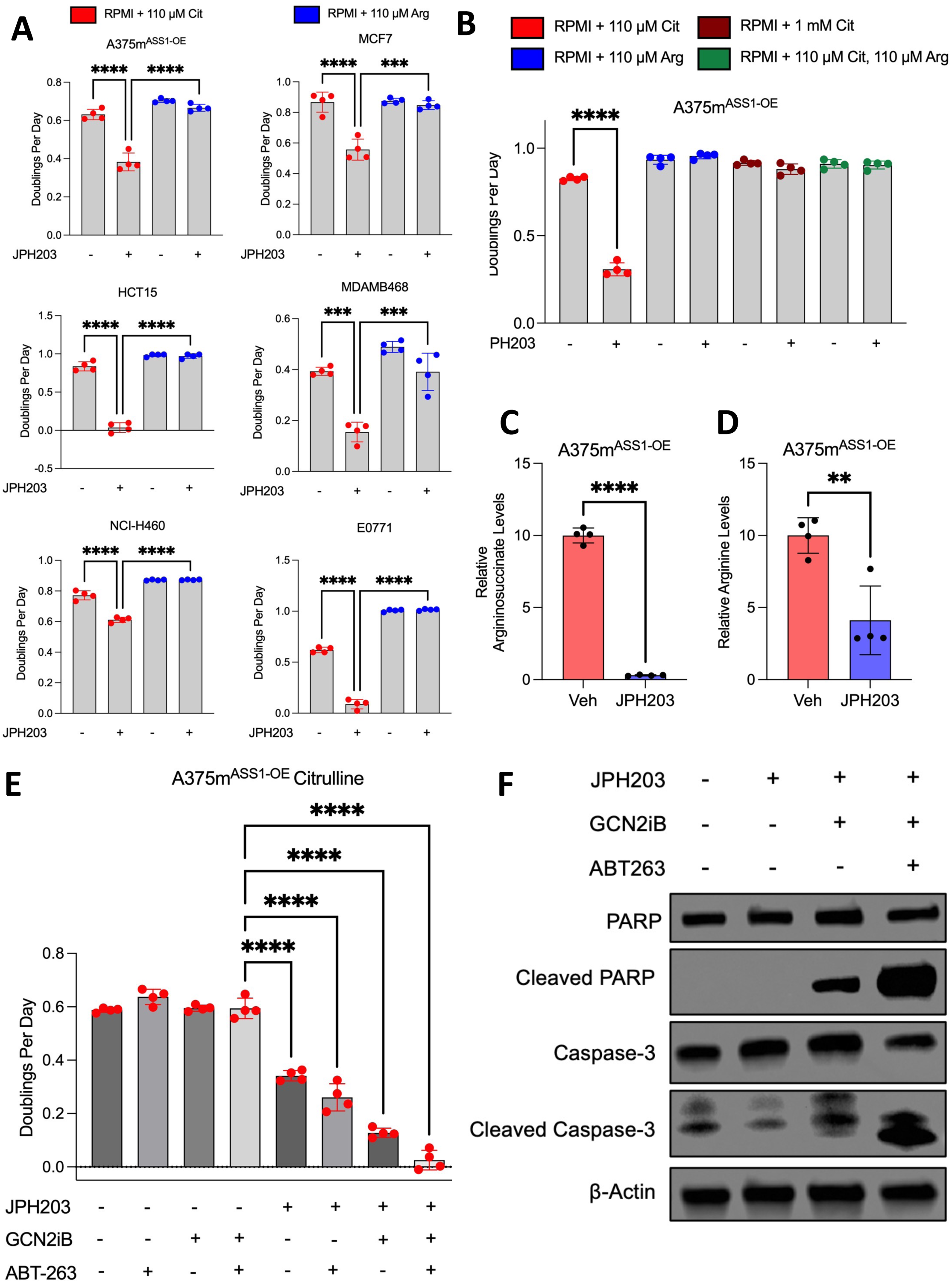
A small molecule inhibitor of SLC7A5 sensitizes cells to arginine deprivation. A: Growth of cancer cell lines cultured in RPMI + Arg, RPMI + Cit, with and without 1 μM JPH203, or vehicle. Data presented after 3 or 4 days of growth, depending on the cell line. Media was refreshed on Day 2. (mean ± SD, n = 4). *** p > .001. **** p < .0001, 2-way ANOVA. B: Rescue of growth in the Cit + JPH203 condition with either 1 mM citrulline or 110 μM Arginine. Media was refreshed on Day 2. (mean ± SD, n = 4). C: Intracellular argininosuccinate levels in A375m^ASS1-OE^ cells with vehicle control or 1 μM JPH203, metabolites extracted after 24 hours of culture in RPMI + 110μM Cit. (mean ± SD, n = 4). **** p < .0001, unpaired, two-tailed t-test. D: Intracellular arginine levels in A375m^ASS1-OE^ cells with vehicle control or 1 μM JPH203, metabolites extracted after 24 hours of culture in RPMI + 110μM Cit. (mean ± SD, n = 4). ** p < .01, unpaired, two-tailed t-test. E: Growth of A375m^ASS1-OE^ cells treated with a combination of inhibitors. JPH203, GCN2iB, and ABT-263 were all dosed at 1μM. Cells grown in RPMI + 110μM Cit. (mean ± SD, n = 4). **** p < .0001, one-way ANOVA. F: Immunoblot of A375m^ASS1-OE^ cells treated for 48 hours in RPMI + 110μM Cit and indicated inhibitors, which were all dosed at 1μM. DMSO was used as a vehicle control in non-drug wells.

Cells deprived of arginine enter senescence, and it was recently reported that senolytic drugs can be paired with arginine starvation to further slow proliferation^7^. We tested whether JPH203 inhibition of citrulline metabolism would yield a similar result. We found that when JPH203 is paired with GCN2iB and the senolytic BCL-2 inhibitor ABT263, proliferation was inhibited when cells are plated in media containing citrulline without arginine (Fig 5E). Importantly, this combination of inhibitors has no effect on cells cultured in arginine (Fig. S5B). We then performed immunoblotting on two apoptotic markers, Cleaved PARP and Cleaved Caspase-3^44–47^. Apoptosis was apparent when JPH203 was combined with GCN2iB and further enhanced by addition of ABT-263 in citrulline dependent growth conditions (Fig 5F). This suggests that cells treated with JPH203 are growth arrested, but not apoptotic until the addition of GCN2iB, which inhibits *ASS1* re-expression, and the senolytic agent ABT-263.

### SLC7A5 regulates citrulline metabolism in an in vivo xenograft model

To examine the effects of SLC7A5 inhibition on citrulline metabolism and tumor growth in an in vivo setting, we injected A375m WT and SLC7A5 KO cells bilaterally into the flanks of 6-week-old female nude mice and tracked tumor growth. Loss of SLC7A5 resulted in significantly reduced tumor growth rate (Fig. 6A, 6B, S6A). To determine levels of citrulline import in these tumors, we injected 0.07g/kg of [1-^13^C]-citrulline intraperitoneally in mice bearing the tumors in Fig. 7A-B and sacrificed the mice 1 hour later. Serum enrichment of [1-^13^C]-citrulline was 34 ± 4% one hour after injection (Fig. 6C). We observed a significant decrease [1-^13^C]-citrulline within KO tumors compared to their WT counterparts in the same mouse (Fig 6D).

**Figure 6:**
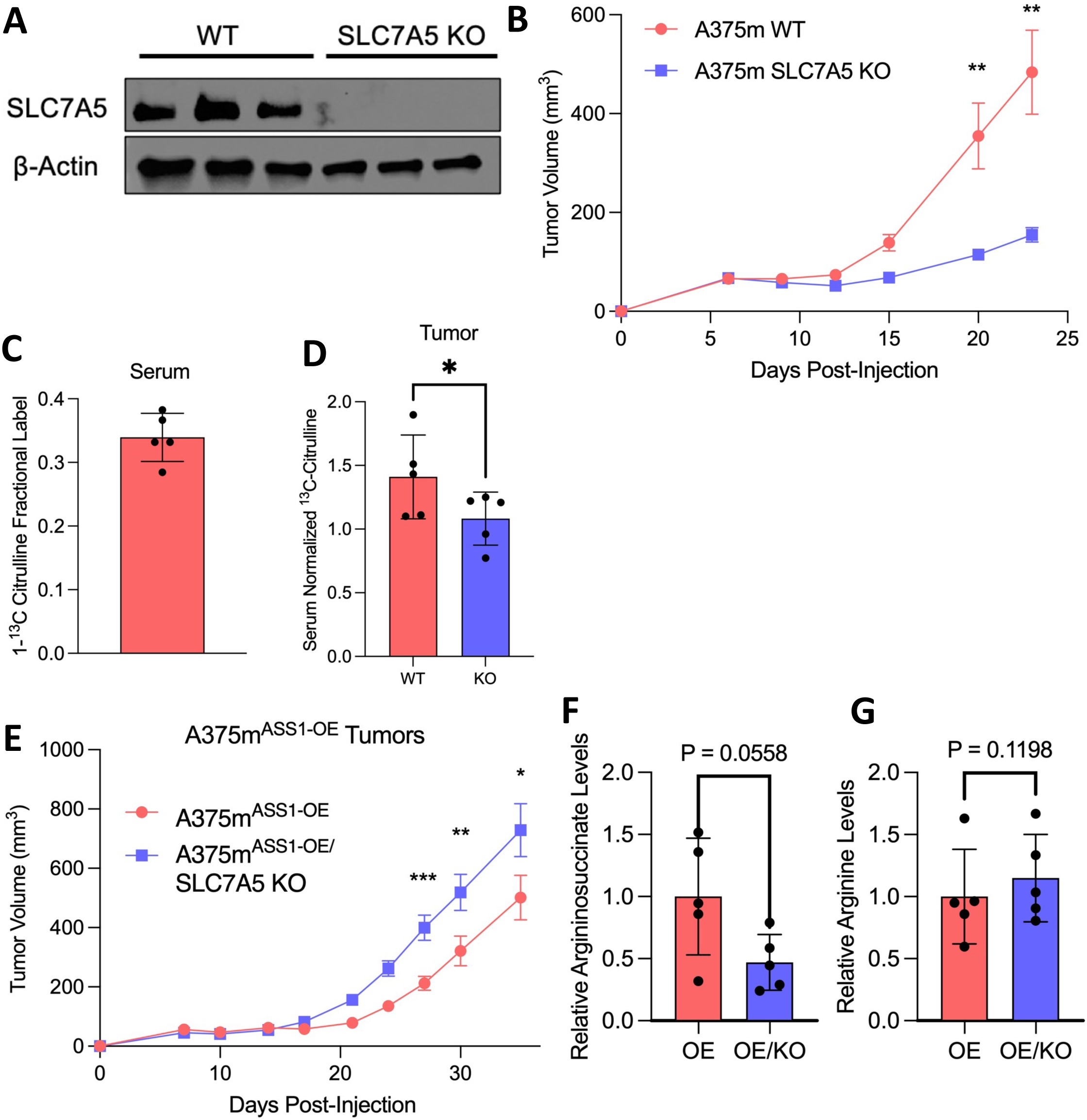
SLC7A5 regulates citrulline metabolism in an in vivo xenograft model. A: Immunoblot of SLC7A5 in tumors originating from A375m xenografts. β-actin used as a loading control. B: Growth of bilateral A375m WT and SLC7A5 KO xenograft tumors. Mice were fed a standard chow diet. (mean ± SEM, n = 10). ** = p < .01, paired, two-tailed t-test. C: Labeling fraction of [1-^13^C]-citrulline in the serum of mice 1 hour after intraperitoneal injection (mean ± SD, n = 5). D: Tumor 1-13C-citrulline uptake normalized to serum citrulline. Mice were injected with ∼83 mM (0.07g/kg) of [1-^13^C]-citrulline. (mean ± SEM, n = 5). * = p < 0.05, paired, two-tailed t-test. E: Growth curves of A375m^ASS1-OE^ WT and SLC7A5 KO xenograft tumors. (mean ± SD, n = 10). * = p < 0.05, ** = p < 0.01, *** = p < 0.001, paired, two-tailed t-test. F: Intra-tumoral argininosuccinate levels in A375m^ASS1-OE^ WT and SLC7A5 KO xenografts. (mean ± SD, n = 5), unpaired, two-tailed t-test. G: Intra-tumoral arginine levels in tumors originating from A375m^ASS1-OE^ WT and SLC7A5 KO xenografts. (mean ± SD, n = 5), unpaired, two-tailed t-test.

Finally, we examined the relationship between ASS1 and SLC7A5 in vivo by utilizing our A375m^ASS1-OE^ and A375m^ASS1-OE^/SLC7A5 KO cells. A375m^ASS1-OE^ tumors grew slower than the parental A375m tumors. These data are in line with suppression of ASS1 in tumors; cancer cells downregulate ASS1 to promote nucleotide production, and forced overexpression of ASS1 exhausts aspartate pools, thereby limiting the growing cells capacity to synthesize necessary nucleotides^21^. Counterintuitively, loss of SLC7A5 in the A375m^ASS1-OE^ tumor rescued tumor growth (Fig. 6E). Consistent with loss of SLC7A5 leading to loss of citrulline uptake, arginosuccinate levels were suppressed in the KO tumors (Fig. 6F). As expected in arginine replete serum, tumor arginine levels however were not changed (Fig. 6G). In summary, these data show that SLC7A5 can interact with ASS1 in vivo to regulate tumor growth, likely via the transport of citrulline.

## Discussion

The uptake of nutrients from the serum serves as a critical control point for metabolic regulation. Differential transport helps underpin cell-type variation in metabolism and contributes to tumor growth. Despite this, the elucidation of many of these transport pathways remains unsolved, although new screening methodologies have now enabled a resurgence in transporter annotation and discovery^31,48–50^. In this study, we asked how a common non-proteinogenic amino acid, citrulline, is used to fuel arginine metabolism. Using a functional genomics screen, we identified the neutral amino acid transporter *SLC7A5* to be an essential gene when cells are dependent upon citrulline uptake for arginine synthesis. Our metabolic analysis revealed that SLC7A5 enables citrulline uptake and conversion into arginine in diverse mammalian cell lines. Small-molecule inhibition of SLC7A5 inhibits cell growth in arginine depleted conditions and, when combined with senolytic agents, leads to apoptosis. Finally, we showed that loss of SLC7A5 results in decreased tumor xenograft growth, which was associated with attenuated citrulline import into tumors.

Diet is a poor source of citrulline in humans, with the most concentrated food source being watermelon (*Citrullus lanatus*)^51^. Data from arterial-venous sampling studies shows that citrulline is largely produced in and exported from intestine enterocytes and synthesized from glutamine and arginine^15,52,53^. A study examining metabolite exchange between organs in pigs show that the leg muscle, spleen, and kidney uptake significant amounts of citrulline, illustrating that localized arginine synthesis is important for protein homeostasis and cell proliferation^54^. Kidney is well-annotated to convert citrulline back into arginine for redistribution in the circulation^55^. Our mouse data matched what was found in pigs, with muscle, spleen and kidney being 3 of the top 4 organs for citrulline import in mice (Fig S6C). Synthesis of arginine appears to be highest in the kidney, heart and brain (Fig. S6D). While the liver has the highest urea cycle activity of any organ^56^, relatively low arginine labelling levels may be related to the high levels of unlabeled arginine present. Our data from circulating citrulline combined with our data from tissue uptake highlights the interplay between citrulline and arginine metabolism in multiple organ systems, far beyond just the liver.

Our discovery of SLC7A5 as a citrulline transporter essential for arginine synthesis places it squarely in the de novo synthesis pathway of arginine. Our results indicate that SLC7A5 is necessary but not sufficient for citrulline uptake and, in Figure 4, we showed that ASS1 expression levels drive relative citrulline consumption fluxes. Outside of hepatocytes, the only way to produce arginine is from citrulline^30^. In tissues, co-expression of ASS1 and SLC7A5 are required for arginine synthesis. In addition to generally supplying arginine for protein homeostasis, SLC7A5 mediated citrulline uptake is important for proliferation. Werner et al. reported that citrulline uptake by SLC7A5 was required for the proliferation of T-cells cultured in arginine depleted conditions^12^. Coupled with this finding, our data suggests that in the proliferative context, SLC7A5 may be the only way to import citrulline required for arginine synthesis.

The role of SLC7A5 as a neutral amino acid transporter has been characterized in detail^34,57^. At physiological levels of leucine and phenylalanine, we show that SLC7A5 is not required for proliferation. Importantly, in our screen and follow-up experiments, leucine and phenylalanine were supplied at standard RPMI concentrations, in excess of physiological values. This was a surprising result and raises the possibility that other lower-affinity transporters for leucine can compensate for the loss of SLC7A5 to sustain proliferation under replete amino acid conditions. These may include other members of the LAT family, such as SLC7A6, SLC7A7 and SLC7A8^58^. In the tumor microenvironment, however, depleted levels of essential amino acids may increase the dependency upon SLC7A5 for growth. In contrast, for citrulline, our data suggests that only very low-affinity transporters outside of SLC7A5 exist to transport this metabolite. The reason for these transporters being low-affinity for citrulline but higher-affinity for leucine and phenylalanine is an avenue for future biochemical investigation. Additionally, since leucine and arginine can both activate the mTOR complex, it remains an open question as to what conditions SLC7A5-dependent transport of leucine and citrulline is required for mTOR activation^59^.

The citrulline transport activity of SLC7A5 is most clearly identifiable when arginine is depleted and/or limiting. This can be accomplished therapeutically by ADI-PEG20, which breaks down serum arginine into citrulline and the ammonium ion. Clinical trials are ongoing to treat *ASS1*-low cancers using ADI-PEG20. In general, these clinical trials have had shown some improvement in interim endpoints, but not durable increases in progression free-survival^60–63^. The limited success of these clinical trials may be due to the fact that treatment with this agent drastically spikes circulating citrulline, and tumors begin to re-express ASS1^24,64^. In this situation, citrulline metabolism enables tumor cells to escape arginine-depleting treatment. Inhibition of SLC7A5 may sensitize tumors to an arginine depletion agent such as ADI-PEG20 and improve therapeutic efficacy. Beyond co-treatment with arginine depleting agents, the efficacy of SLC7A5 agents may reveal tumors with limited arginine availability in the TME^65,66^. Numerous studies have shown that JPH203 treatment slows tumor growth in xenograft models^67–70^. Interestingly, these tumors have not been metabolically profiled to examine which of the amino acids imported by SLC7A5 drives this reduction in tumor burden. Given findings in T-cells, SLC7A5 inhibition may have deleterious roles for T-cell antitumor activation in some contexts^71^. Overall, the finding that SLC7A5 is required for proliferation in arginine-low environments identifies tumor contexts that may be particularly amenable to treatment with these agents.

## Limitations

Metabolomics and isotope tracing suggest a role of SLC7A5 as the main citrulline transporter in mammalian cells. Cell lines do not fully recapitulate in vivo variation, and it is possible that other transporters besides SLC7A5 perform this function in vivo. The data from the CRISPR screen performed in this study is limited by the choice of cell line and media. In primary cells, there may be additional high-affinity transporters that can fully rescue citrulline import and arginine-deficient growth when SLC7A5 is absent. Additionally, the mice used for tumor modeling in this study were immunodeficient. Arginine has been suggested to play a significant role in T-cell metabolism^10^. Further studies using immunocompetent mouse models will be required to understand the therapeutic feasibility of inhibiting citrulline uptake in the tumor microenvironment.

## Acknowledgements

We thank Kivanc Birsoy for the Human Metabolism CRISPR Knockout Library. We thank Adam Hughes for his input on the experimental strategy and close reading of the manuscript. Research reported in this publication utilized the High-Throughput Genomics and Cancer Bioinformatics Shared Resource at Huntsman Cancer Institute at the University of Utah. Research reported in this publication also utilized the Preclinical Research Shared Resource at Huntsman Cancer Institute at the University of Utah. Both are supported by the National Cancer Institute of the National Institutes of Health under Award Number P30CA042014. Additionally, we thank the Flow Cytometry Core at the University of Utah, supported by the Office of The Director of the National Institutes of Health under Award Number S10OD026959. The content is solely the responsibility of the authors and does not necessarily represent the official views of the NIH. G.S.D. was supported by funding from the NIH (R00 CA215307), The American Cancer Society (DBG-23-1037804-01-TBE) and the Nuclear Control of Cell Growth and Differentiation Research Group of the Huntsman Cancer Institute of the University of Utah.

## Author Contributions

Conceptualization, K.N.D., A.J.B. and G.S.D.; methodology, K.N.D and G.S.D.; analysis, K.N.D and G.S.D.; investigation, K.N.D, A.Be., A. Bo., A.J.B.; writing and editing, K.N.D., A.J.B, J.R. and G.S.D.; funding acquisition G.S.D.

## Declaration of Interests

The authors declare no conflicts of interest.

## Methods

### Cell Lines

A375m, MCF7, HCT15, HCT116, EO771, NCI-H460, LN229 and MDA-MB-468 cell lines were maintained at 37°C and 5% CO_2_ and tested regularly for mycoplasma contamination using a mycoplasma detection kit (Applied Biological Materials, G238). Cells were cultured in either RPMI (Thermo, 11875) or DMEM (Thermo, 11965). The media was supplemented with 10% fetal bovine serum (FBS). (Thermo, 10437028) and 100 U/mL penicillin/100 µg/mL streptomycin (Thermo, 15140122). MDA-MB-468 cells were supplemented with 5% sodium pyruvate (Thermo, 11360070).

### Lentivirus Production and Transfection to Make Overexpressing Cell Lines

7.5×10^5^ HEK293-FT cells were plated in each well of a 6-well dish coated with 50 μg/mL of Poly-D-Lysine (Thermo, A3890401). ASS1 and SLC7A5 cDNA in the vectors plX304 (Addgene #25890) and plX307 (#41392), respectively, along with packaging vectors psPAX2 (#12260) and PMD2.G (#12259) were transfected into the HEK293-FT cells using XTremeGene 9 (Sigma Aldrich, 6365779001) (A375m) or Lipofectamine 3000 (Thermo, L3000008) (MCF7). Media was changed 6 to 12 hours after transfection. The virus-containing supernatant was collected 36 to 48 hours after the media change, spun at 400 x g for 5 minutes at room temp, and passed through a .45 μm filter to eliminate cells.

For infection, we infected 50,000 cells in 6-well tissue culture plates with 1mL media containing 10 μg/mL polybrene and either 500, 300, 150, 30, 15 or 0 μL of virus. 2 days after infection, the virus-containing media was removed and changed to media containing either 1 μg/mL blasticidin (A375m, ASS1 cDNA), or 1 μg/mL puromycin (A375m, SLC7A5 cDNA). The cells that survived blasticidin or puromycin treatment were expanded, overexpression of protein was validated using western blotting, and used for downstream experiments.

### Generation of CRISPR Knockouts

To make CRISPR knockouts, we followed the protocol in Ran, 2013^26^. Briefly, sgRNAs (oligonucleotide sequences in the table elsewhere) were cloned into pSpCas9(BB)-2A-Puro (Addgene #62988) or pSpCas9(BB)-2A-GFP (#48138). The resulting plasmid was then transfected into cells using Lipofectamine 3000. After 2-3 days, cells were either sorted by FACS for GFP, or selected for puromycin. After selection, cells were single-cell sorted with a flow cytometer into the wells of a 96-well plates containing 100 μL of RPMI and 15% FBS. Cells were grown until sufficient size, and then expanded so that the loss of the target protein could be validated via western blotting, and then used for downstream experiments.

### Cell Lysate Preparation and Western Blotting

Cells were plated in 6 well plates and allowed to grow for 1-4 days (depending on the experiment) at 37°C and 5% CO_2_. The cells were washed with cold PBS and lysed in RIPA buffer (50 mM Tris, 150 mM NaCl, 0.1% SDS, 0.5% sodium deoxycholate, 1% NP-40) supplemented with 1 mM phenylmethylsulfonyl fluoride (PMSF, Cell Signaling 8553S), 1 mM sodium orthovanadate (Sigma-Aldrich, S6508), and 1% protease inhibitor cocktail (Sigma, P2714). The lysing occurred by scraping the cells off the plates with a cell scraper. Lysate was then agitated on ice for 15-30 minutes and then briefly sonicated (Branson Ultrasonics) and centrifuged at 16,000 x g for 15 minutes at 4C. Supernatant containing protein was then placed in a fresh tube. Protein concentration was quantified using the Pierce BCA Protein Assay Kit (Thermo, 23225). Samples were mixed with 3X Laemmli Sample Buffer (BioRad, 1610747) and 2-mercaptoethanol to a final concentration of 3.33% and incubated at 80-90°C for 10 min. Depending on the protein target, 1-20 μg of total protein was resolved on a 4-20% polyacrylamide gel (BioRad, 4561096) at 100 V for 60-90 minutes. Gels were blotted on 0.2 µm nitrocellulose membranes (BioRad, 1704270) via the Trans-Blot Turbo Transfer System (BioRad, 1704150) according to standard procedure at 25 V for 30 min. After blocking with 5% non-fat milk (Sigma, M7409)/Tris-buffered saline with 0.05% Tween 20 (TBS-T) for 1 hr, the membrane was washed with TBS-T and incubated overnight in 5% bovine serum albumin (Sigma, A6003)/TBS-T with 1:1000 of the primary antibody (see the materials table for a list). The membrane was then washed with TBS-T and incubated with 1:10000 goat anti-rabbit or rabbit anti-mouse poly-HRP secondary antibody (Thermo, 32260) and 1:5000 anti-β-actin-peroxidase antibody (Sigma, A3854) in TBS-T for 1 hr. The membrane was then washed with TBS-T and chemiluminescence was assessed with the BioRad ChemiDoc MP Imaging System (BioRad, 12003154).

### Proliferation Assays

Depending on the cell line, between 1500 and 5000 cells were plated in each well of a 96 well plate. For growth experiments not involving drugs, cells were plated in their indicated media. For experiments involving JPH203, GCN2iB or ABT-263, cells were plated in RPMI + 110 μM Arginine, and changed to experimental media the next day. Media was changed every 2 days. At indicated time points, 10 μL per 100 μL of media of .03% resazurin salt (Sigma, R7017) solution was added and incubated for 2 hours. After this incubation period, cell growth was read as fluorescence intensity using a multi-mode plate reader (BioTek) and expressed as relative growth.

### Metabolite Extraction in vitro

Cells were grown in 6 cm tissue culture dishes for 16-48 hours, depending on the experiment. For amino acid labeling experiments, media was changed at indicated times before harvesting. Cell plates were washed with ice cold PBS 2x and then 80:20 methanol:water was added at 60x of the PCV (packed cell volume) (MidSci, TP87005) count. The resulting mixture was incubated on dry ice, scraped, collected into a microfuge tube, vortexed, rested on dry ice for 5 minutes and centrifuged at 16000 × g for 10 min. Supernatant was placed into a fresh tube which was then centrifuged again at 16000 x for 10 min. The supernatant was placed in an MS tube (Agilent 5188-2788) for downstream analysis.

### Creation of Custom Media

Media involving citrulline, leucine, or arginine manipulations were made from a source stock containing all components of RPMI without leucine, lysine, and arginine (US Biologicals, R8999-03A). Media involving phenylalanine manipulations originated with a -glucose and -amino acid stock (US Biologicals, R9010-02). Experimental media was then made by adding back in amino acids at concentrations indicated in individual experimental protocols (pH 7.4).

### RNA isolation and Quantitative Real-Time PCR

Cells were plated on 6-well plates and allowed to grow for 1-4 days (depending on the experimental setup) at 37°C and 5% CO_2_. Cells were washed with PBS and RNA was isolated using the Qiagen RNeasy Mini Kit (Qiagen, 74104) according to the manufacturer’s specifications. cDNA was then generated using QuantaBio qScript cDNA SuperMix (QuantaBio 101414-102). qPCR master mixes were prepared consisting of 2.5% 100 µM forward primer, 2.5% 100 µM reverse primer, and 62.5% PowerTrack SYBR Green Master Mix (Thermo, A46109) in RNase-free water. Master mixes were combined 4:1 with the cDNA reactions and plated in duplicate. qPCR was performed using the LC480 PCR Lightcycler (Roche, 05015278001) using the “Mono Color Hydrolysis Probe/UPL probe” detection format. The temperature cycle consisted of an initial 2 min period at 95°C and 40 cycles of 95°C for 15 sec and 60°C for 50 sec set to single acquisition mode. The housekeeping gene *RPS2* was used as an internal control for cDNA quantification and normalization of the amplified products.

### Metabolite Extraction from Serum

4 μL of serum was added to 68 μL 100% methanol, vortexed, and put on dry ice for at least 5 minutes. This mixture was then centrifuged for 10 minutes at 16,000 x g at 4C. Supernatant was placed into a fresh tube and mixed 1:1 with 80% methanol. The sample was vortexed and centrifuged again for 10 minutes at 16,000 x g at 4C. 100 μL of supernatant was transferred toan MS tube for LC-MS analysis.

### Metabolite Extraction from Tissue

30-40 mg sections of snap frozen mouse tissue were transferred to pre-chilled Safe-Lock tubes (Eppendorf, 022363352) containing a cold 5/16 in. diameter stainless steel ball (Grainger, 4RJL8). The tissue was disrupted by shaking at 25 Hz for 30 sec under liquid nitrogen using the Retsch CryoMill (Retsch, 20.749.0001). 15 μL per mg of tissue of a polar metabolite extraction solution containing 40:40:20 Acetonitrile:Methanol:Water and .1% Formic Acid was added to homogenized tissue Samples were briefly vortexed before neutralizing with 8 mL of 15% ammonium bicarbonate per 100 mL of extraction solvent. Samples were vortexed again and centrifuged at 16,000 x g for 10 minutes at 4C. Supernatant was put into a fresh tube. 40:40:20 ACN:MeOH:H2O with no formic acid was added to the original tube, vortexed, centrifuged at 16,000 x g for 10 minutes at 4C. Supernatant was added to the same fresh tube mentioned above. 50% chloroform was added to the fresh tube and vortexed to induce phase separation. Samples were again vortexed and centrifuged at 16,000 x g for 10 minutes at 4C. Supernatant (∼400 μL) was placed into a fresh tube and centrifuged at 16,000 x for 10 minutes at 4C for the final time. Supernatant (∼200 μL) was placed in an MS tube for LC-MS analysis.

### CRISPR/Cas9 Functional Genomic Screens

Metabolism-scale functional screens using CRISPR were performed as described in previous work^31,32^. Briefly, 2,989 genes encoding metabolic enzymes, some transcription factors, and small molecular transporters were targeted with a total of 23,777 sgRNAs and 50 controls targeting intergenic regions. After cloning into lentiCRISPR-v2 puro (Addgene #982990), the pooled plasmid library was used to produce lentivirus-containing supernatants. The optimal volume of lentiviral supernatants to use for the screen was determined by infection of target cells at a range of virus and counting the number of puromycin-resistant cells after 3 days of selection. For the screens, 100×10^6^ target cells were infected at an MOI of ∼0.3 and selected with 1 μg/mL puromycin. An initial pool of ∼30×10^6^ cells were harvested for genomic DNA extraction at the beginning of the screen. Blasticidin was present in the media at a concentration of 1 μg/mL throughout the duration of the screen, due to the ASS1 overexpressor construct. 24×10^6^ cells at a concentration of 4.8×10^6^ per 15cm plate were subjected to each experimental condition and passaged every 2 days until 14 cumulative population doublings were reached. On the final day of screening, cells were harvested for genomic DNA extraction. Genomic DNA was extracted using a DNeasy Blood & Tissue Kit (Qiagen) and amplification of sgRNA inserts was performed via PCR using barcoded primers for each condition. PCR amplicons were purified and sequenced on a NovaSeq (Illumina). Fastq files were analyzed using MAGeCK software. Gene score is defined as the median log_2_ fold-change in the abundance of all sgRNAs targeting a particular gene between the initial and final populations.

### LC-MS Methodology

Extracted aqueous and polar metabolites were analyzed by LC-MS using a Vanquish HPLC system (Thermo Fisher Scientific) and a QExactive Plus Orbitrap mass spectrometer (Thermo Fisher Scientific). For aqueous phase polar metabolites, separation was achieved by hydrophilic interaction liquid chromatography (HILIC) performed on an Atlantis Premier BEH Z-HILIC column or a Waters BEH HILIC column. For the Z-HILIC column, the specifications were as follows: (2.1 mm X 50 mm, 1.7 µM particular size, 95 Å pore size, Waters Co., 186009978) run with a gradient of solvent A (10 mM ammonium acetate in 100% water, pH 9.2) and solvent B (100% acetonitrile) at a constant flow rate of 350 µL/min. The gradient function was: 0 min, 95% B; 10.4 min, 45% B; 11.5 min, 45% B; 11.6 min, 95% B; 15 min, 95% B. For the HILIC column, the specifications are as follows: (2.5 µm particular size, 2.1 mm X 150 mm, 230 Å pore size, Waters Co., 186006724) run with a gradient of solvent A (10 mM ammonium acetate in 100% water, pH 9.2) and solvent B (100% acetonitrile) at a constant flow rate of 350 µL/min. The gradient function was: 0 min, 90% B; 3 min, 75% B; 9 min, 70% B; 10 min, 50% B; 13 min, 25% B. For both columns, the autosampler temperature was 4°C, column temperature was 30°C, and injection volume was 3 µL. Samples were injected into the mass spectrometer by electrospray ionization operating in either negative or positive ion mode. Samples were analyzed using a full scan method with a resolving power of 70,000 at m/z of 200 and range of 74 – 1110 m/z. Full scan data were analyzed using the Maven software package with specific peaks assigned based on exact mass and comparison with known standards^72^. Extracted peak intensities were corrected for natural isotopic abundance using the R package AccuCor^73^.

### Citrulline Uptake and Import Measurements

For the short-term uptake measurements, cells were plated in 6cm dishes overnight. 15 minutes before metabolite extraction, media was changed to RPMI containing 40 μM [1-^13^C]-Citrulline (Cambridge Isotopes). Then, metabolites were extracted from cells as described previously and run on the LC-MS.

For long-term experiments, cells were plated and media was taken, metabolites extracted, and run on the LC-MS at the time of plating (time 0) and 48 hours later. To normalize to cell count, packed cell volume (PCV) measurements were taken for each cell line at 0, 24, and 48 hours after plating. The equation used to calculate nmol Citrulline or Arginine consumed per hr/μL of cells is: 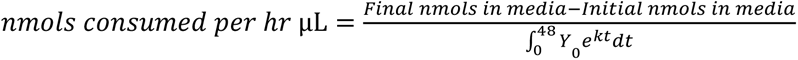

### In vivo Isotope Tracing

Mice with catheters pre-placed in the jugular vein were purchased from Charles River. [1-^13^C]-citrulline (10 mM, 0.1 μl g^−1^ min^−1^), or [^13^C_6_]-arginine (20 mM, 0.1 μl g^−1^ min^−1^) (Cambridge Isotopes) were infused into 3-hour fasted mice for 3 hours. For non-terminal infusions, a bolus at the rate of 1.6 μl g^−1^ min^−1^ for one minute at the beginning of the infusion. Blood samples were collected from the tail vein to measure the enrichment of infused isotopes. For experiments measuring intra-organ labeling, mice were euthanized after the 3-hour infusion by cervical dislocation and various organs were quickly snap frozen in liquid nitrogen using a pre-cooled Wollenberger clamp. Organ labeling was normalized to serum citrulline label. For the 1-hour citrulline injection in Figure 7, mice were fasted overnight and ∼83 mM (0.07g/kg) of [1-^13^C]-Citrulline was injected intraperitoneally. mice were taken down 1 hour after the citrulline injection.

### Mouse Xenografts

All animal studies were approved and conducted under the supervision of the University of Utah Institutional Animal Care and Use Committee. For mice injected with A375m^ASS1-OE^ cells, it was done so at a concentration of 4×10^5^ cells in 100 μL of Matrigel, and cells were injected into the flank. For A375m parental cells, the concentration was 3×10^5^ cells in 100 μL of Matrigel. These mice were given a standard chow diet. For these experiments, WT and SLC7A5 KO genotypes were compared. WT and KO cells were implanted bilaterally on the same animal in growth media. Tumor growth measurements were taken biweekly. Animals were euthanized when the largest tumors reached approximately 600-1000 mm^3^ or if they displayed any signs of distress or morbidity.

---

**Supplemental Figure 1.**
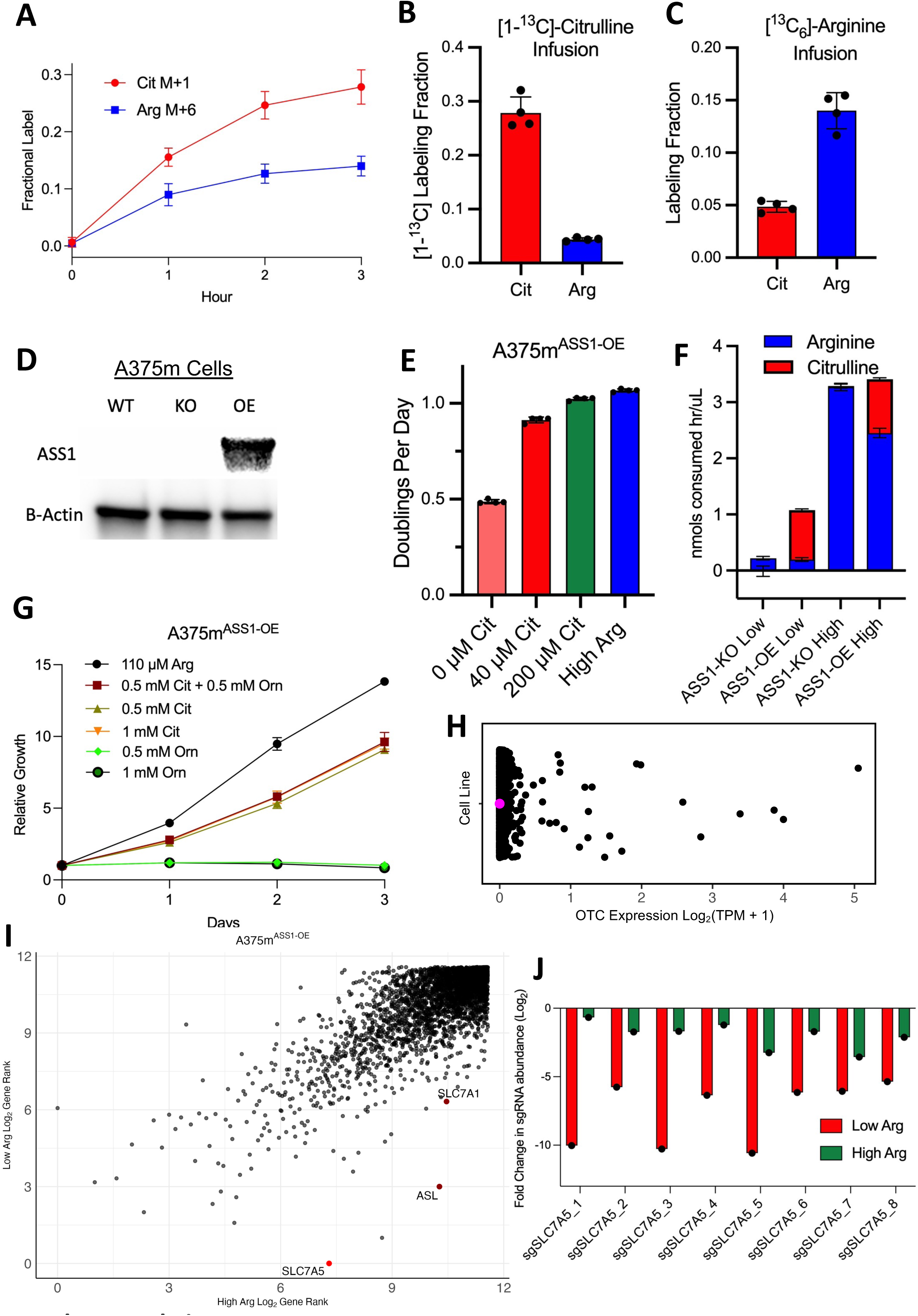
A: Fractional label of serum M+1 citrulline or M+6 arginine at indicated time points during a 3-hour infusion with 10 mM [1-^13^C]-Citrulline or 20 mM [^13^C_6_]-Arginine. B: [1-^13^C]-Citrulline or [1-^13^C]-Arginine labeling in the serum after 3 hours of a [1-^13^C]-Citrulline infusion. C: Fully-carbon labelled [^13^C-5]-Citrulline or [^13^C_6_]-Arginine labeling in the serum after 3 hours of a [^13^C_6_]-Arginine infusion. D: Immunoblot of A375m cells containing WT ASS1, ASS1 knocked out by CRISPR (KO), or ASS1 overexpressed (OE) by a lentiviral construct. β-actin is used as a loading control. E: Growth curve of A375m^ASS1-OE^ cells grown in various concentrations of citrulline and arginine in RPMI media without arginine. (mean ± SD, n = 4). F: Consumption of arginine or citrulline from the media in A375m^ASS1-KO^ or A375m^ASS1-OE^ cells in either Low Arg (Low) or High Arg (High). (mean ± SD, n = 3). G: A375m^ASS1-OE^ cells grown over 3 days in various amounts of arginine (Arg), citrulline (Cit) and ornithine (Orn). (mean ± SD, n = 4). H: Transcript expression data across Log_2_(TPM+1) of OTC across 1,489 cell lines in the Cancer Cell Line Encyclopedia (CCLE). Expression of OTC in A375 cells is highlighted in magenta. I: Essentiality ranks for each gene in both conditions, ranked 1 through 2,989. Genes are ranked by their essentiality score from most essential (#1) to least essential (#2989). Data presented as Log_2_ of the rank. J: Changes in abundance in the individual SLC7A5 sgRNAs in Low Arg and High Arg.

**Supplemental Figure 2.**
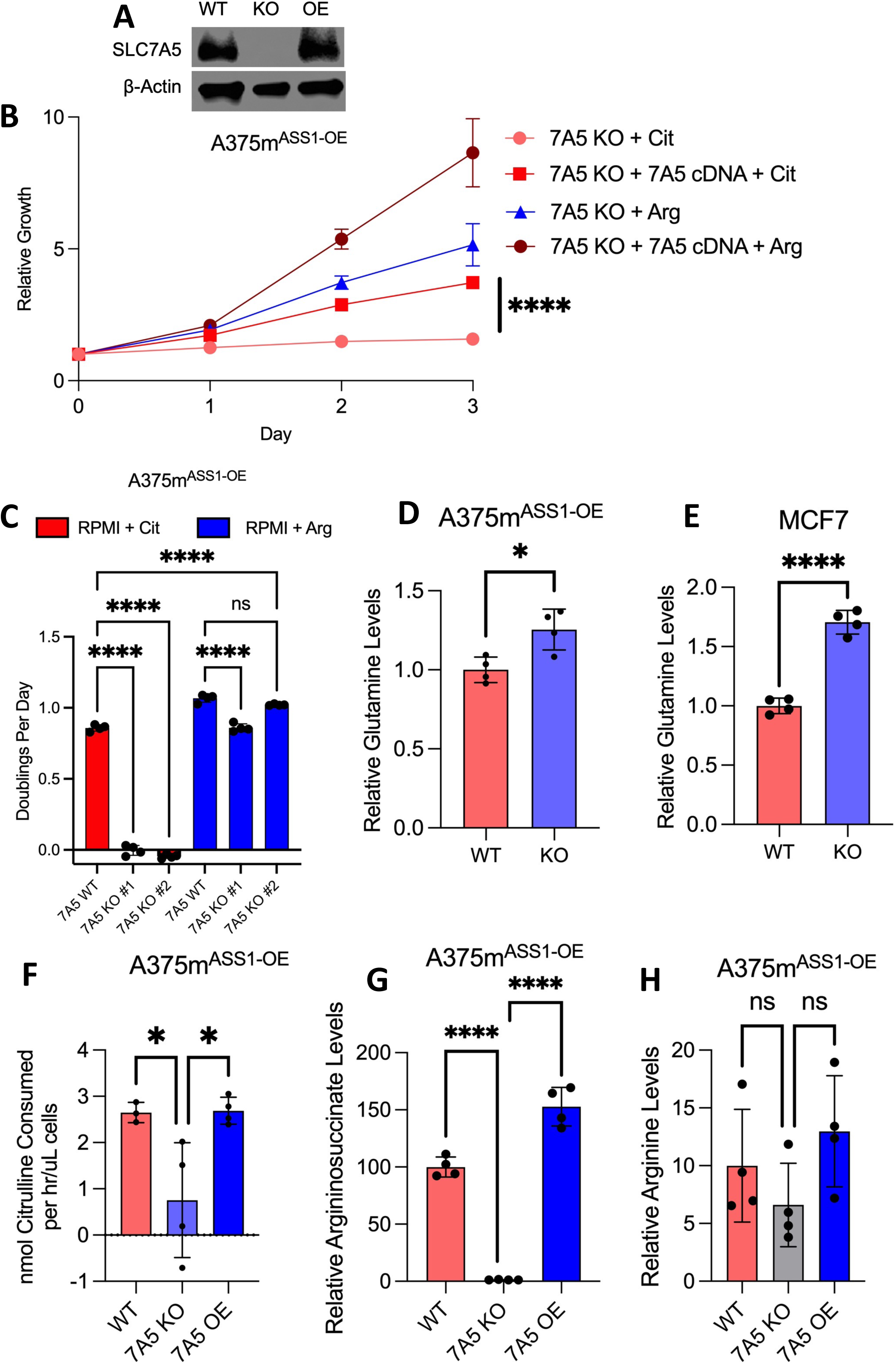
A: Immunoblot of SLC7A5 re-expression in A375m^ASS1-OE^ SLC7A5 KO cells. β-actin is used as a loading control. B: Growth curve of A375m^ASS1-OE^ cells with SLC7A5 knocked out by CRISPR (7A5 KO), or 7A5 KO cells with a lentiviral SLC7A5 cDNA (7A5 cDNA) construct present. Cells grown in either RPMI + 110 μM Arg or RPMI + 110 μM Cit. (mean ± SD, n = 4). **** p < .0001, unpaired, two-tailed t-test at the last day timepoint. C: A375m^ASS1-OE^ WT and SLC7A5 KO cells were grown in indicated media. Media was refreshed on day 2. Data represented as doublings per day. n.s = not significant, (mean ± SD, n = 4), **** p < .0001, one-way ANOVA. D: Intracellular glutamine levels in A375m^ASS1-OE^ cells, metabolites extracted after 24 hours of culture in RPMI + 110 μM Cit. (mean ± SD, n = 4). * p < .05, unpaired, two-tailed t-test. E: Intracellular glutamine levels in MCF7 cells, metabolites extracted after 24 hours of culture in RPMI + 110 μM Cit. (mean ± SD, n = 4). **** = p < .0001, unpaired, two-tailed t-test. F: Citrulline consumption from the media with A375m^ASS1-OE^ cell line with SLC7A5 WT, SLC7A5 KO, and SLC7A5 cDNA (7A5 OE) over a 48-hour period. Media: RPMI + 110 μM Arg + 110 μM Cit. (mean ± SD, n = 4). * p < .05, unpaired, two-tailed t-test. G: Intracellular argininosuccinate in A375m^ASS1-OE^ cells, SLC7A5 WT, KO and OE. Metabolites extracted after 24 hours of culture in RPMI + 110μM Cit. (mean ± SD, n = 4). **** p < .0001, one-way ANOVA. H: Intracellular arginine levels in A375m^ASS1-OE^ cells, SLC7A5 WT, KO and OE. Metabolites extracted after 24 hours of culture in RPMI + 110μM Cit. (mean ± SD, n = 4). n.s not significant, one-way ANOVA.

**Supplemental Figure 3.**
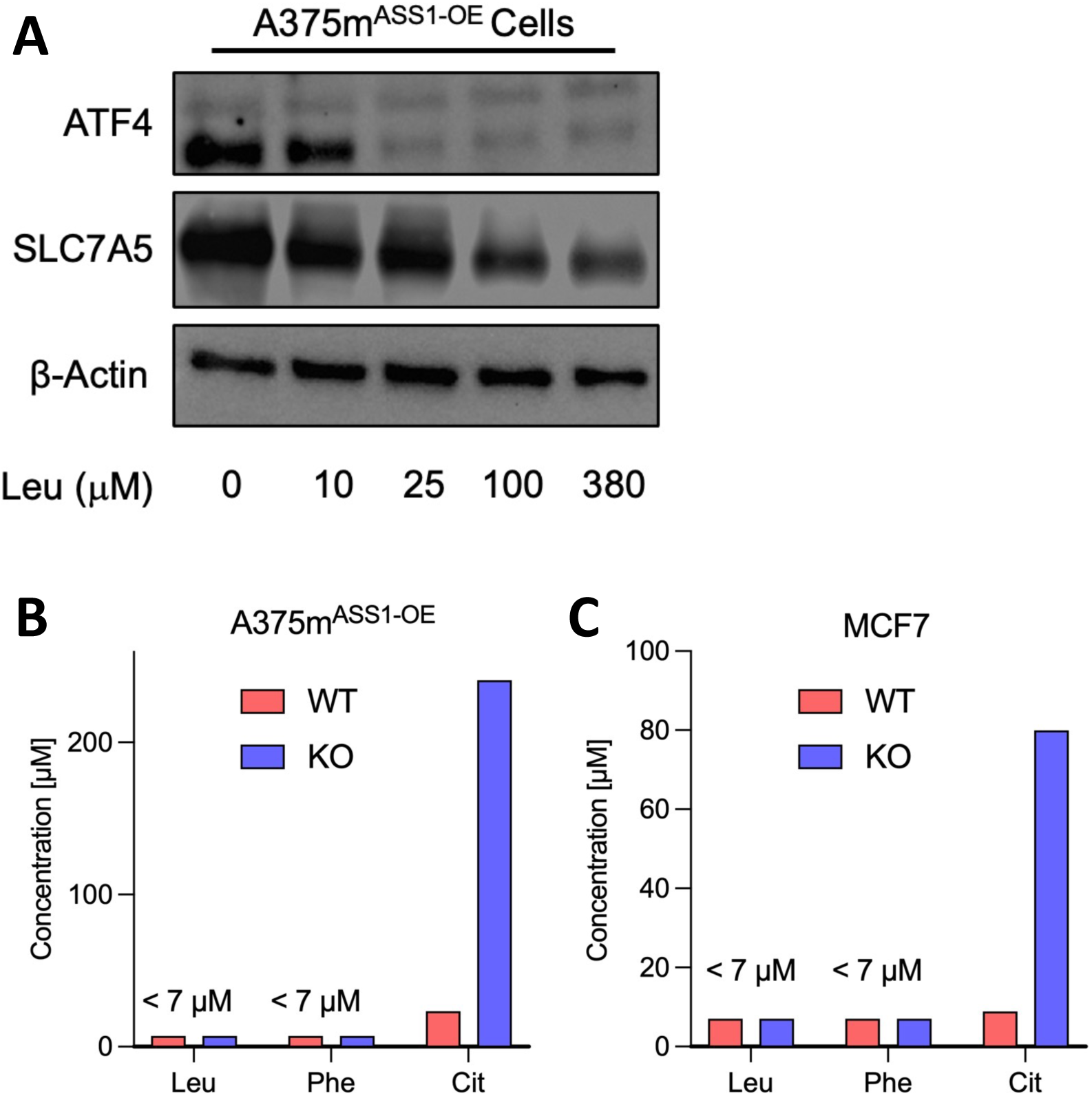
A: Immunoblot of ATF4 and SLC7A5 in RPMI + variable amounts of leucine, cultured for 48 hours before protein extraction. β-actin is used as a loading control. B: Concentration of indicated amino acid required to support the growth rate of half of the peak growth rate of the respective metabolite and genotype combination. Data extrapolated from the best-fit line in Fig. 3, where the concentration (X) was solved for given a known Y (half of the doublings per day at the highest concentration of that amino acid). A375m^ASS1-OE^ cells. C: Concentration of indicated amino acid required to support the growth rate of half of the peak growth rate of the respective metabolite and genotype combination. Data extrapolated from the best-fit line in Fig. 3, where the concentration (X) was solved for given a known Y (half of the doublings per day at the highest concentration of that amino acid). MCF7 cells.

**Supplemental Figure 4.**
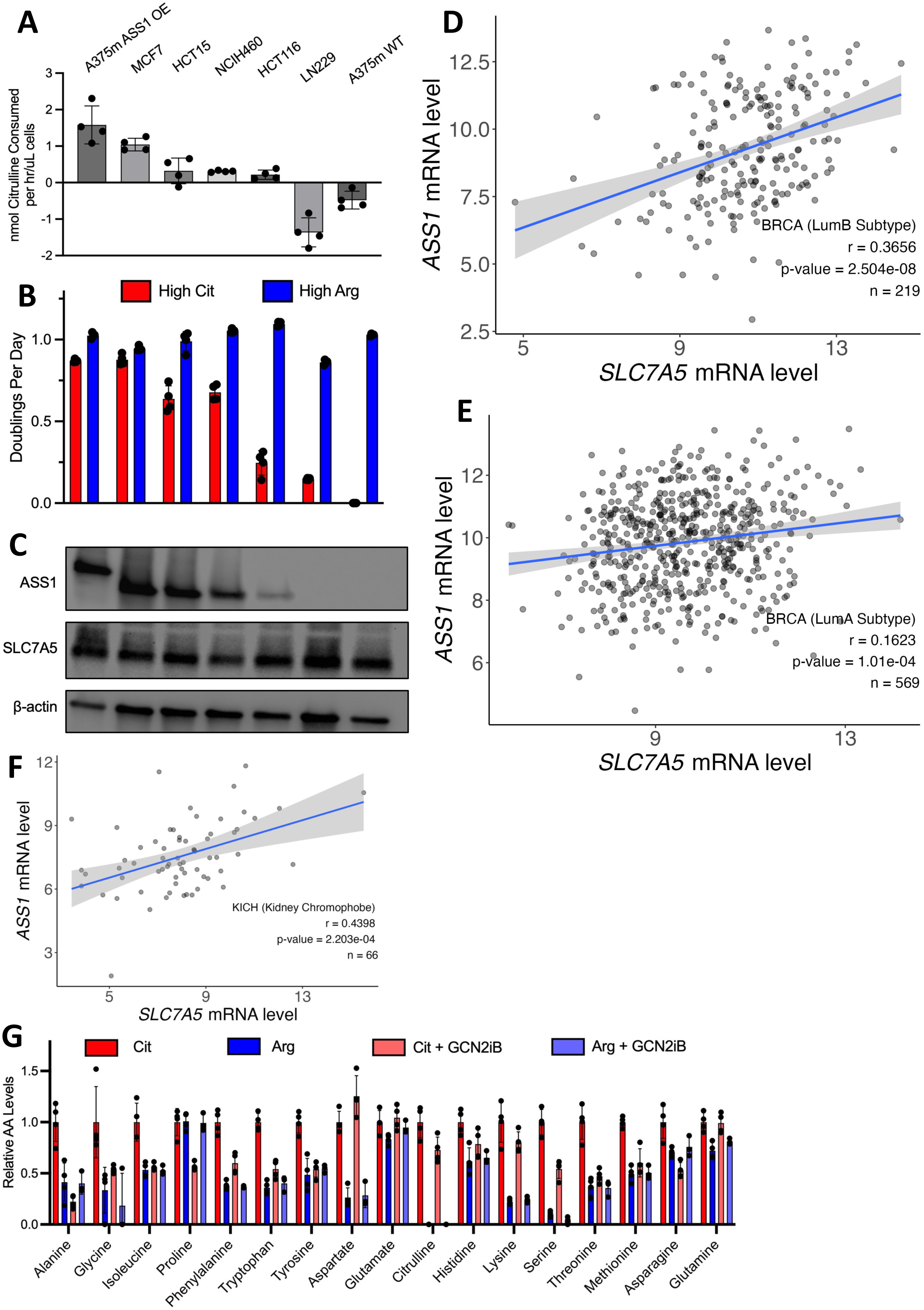
A: Citrulline consumption from the media in a panel of cell lines over a 48-hour period. Media: RPMI + 110 μM Arg + 110 μM Cit. (mean ± SD, n = 4). The cell line labels in panel A correspond to the groups directly below in B and C. B: Growth in RPMI + 110μM Cit (red) or RPMI + 110μM Arg (blue) in a doublings per day format. Growth data is presented after either 3 or 4 days of growth, depending on the cell line. Media was refreshed on day 2. (mean ± SD, n = 4). C: Immunoblot of a panel of proliferating human cell lines. ASS1 and SLC7A5 were blotted for, with β-actin as a loading control. D: Correlation between mRNA levels of *SLC7A5* versus *ASS1* in Luminal B (LumB) mRNA subtypes of breast cancer (BRCA) tumor transcripts data taken from TCGA. The gray area around the linear regression line indicates the 95% confidence interval (CI). E: Correlation between mRNA levels of *SLC7A5* versus *ASS1* in Luminal A (LumA) mRNA subtypes of breast cancer (BRCA) tumor transcripts data taken from TCGA. The gray area around the linear regression line indicates the 95% confidence interval (CI). F: Correlation between mRNA levels of *SLC7A5* versus *ASS1* in kidney chromophobe (KICH) tumor transcript data taken from TCGA. The gray area around the linear regression line indicates the 95% confidence interval (CI). G: Relative amino acid levels of cells plated for 24 hours in either RPMI + 110 μM Cit (Cit, red), RPMI + 110 μM Arg (Arg, blue), RPMI + 110 μM Cit + 1 μM GCN2iB (Cit + GCN2iB, pink) or RPMI + 110 μM Arg + 1 μM GCN2iB (Arg + GCN2iB, periwinkle).

**Supplemental Figure 5.**
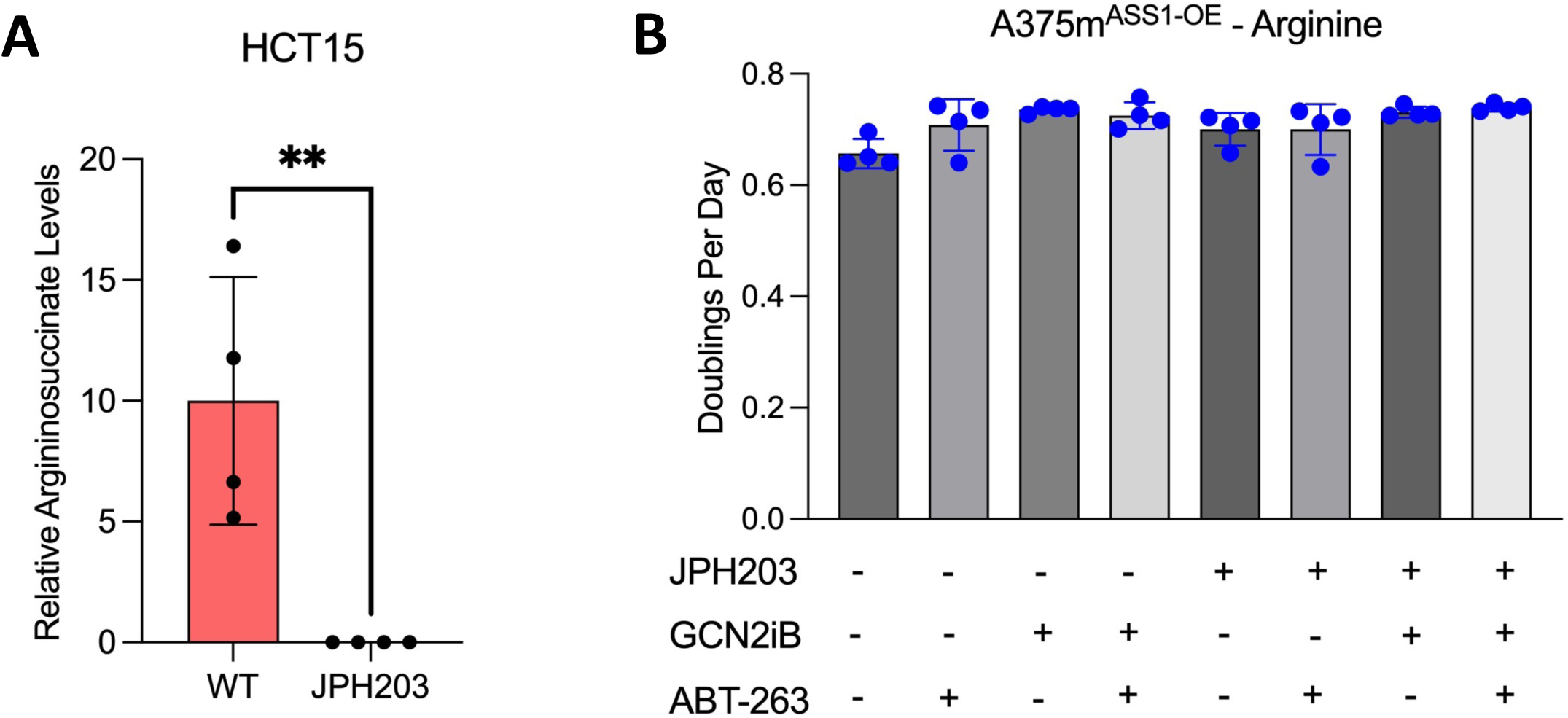
A: Intracellular argininosuccinate levels in HCT15 cells with vehicle control or 1 μM JPH203, metabolites extracted after 24 hours of culture in RPMI + 110μM Cit. Argininosuccinate was not detected by LC-MS in the JPH203 condition. (mean ± SD, n = 4). ** p < .01, unpaired, two-tailed t-test. B: Growth of A375m^ASS1-OE^ cells treated with a combination of inhibitors. JPH203, GCN2iB, and ABT-263 were all dosed at 1 μM. Cells grown in RPMI + 110 μM Arg. (mean ± SD, n = 4).

**Supplemental Figure 6.**
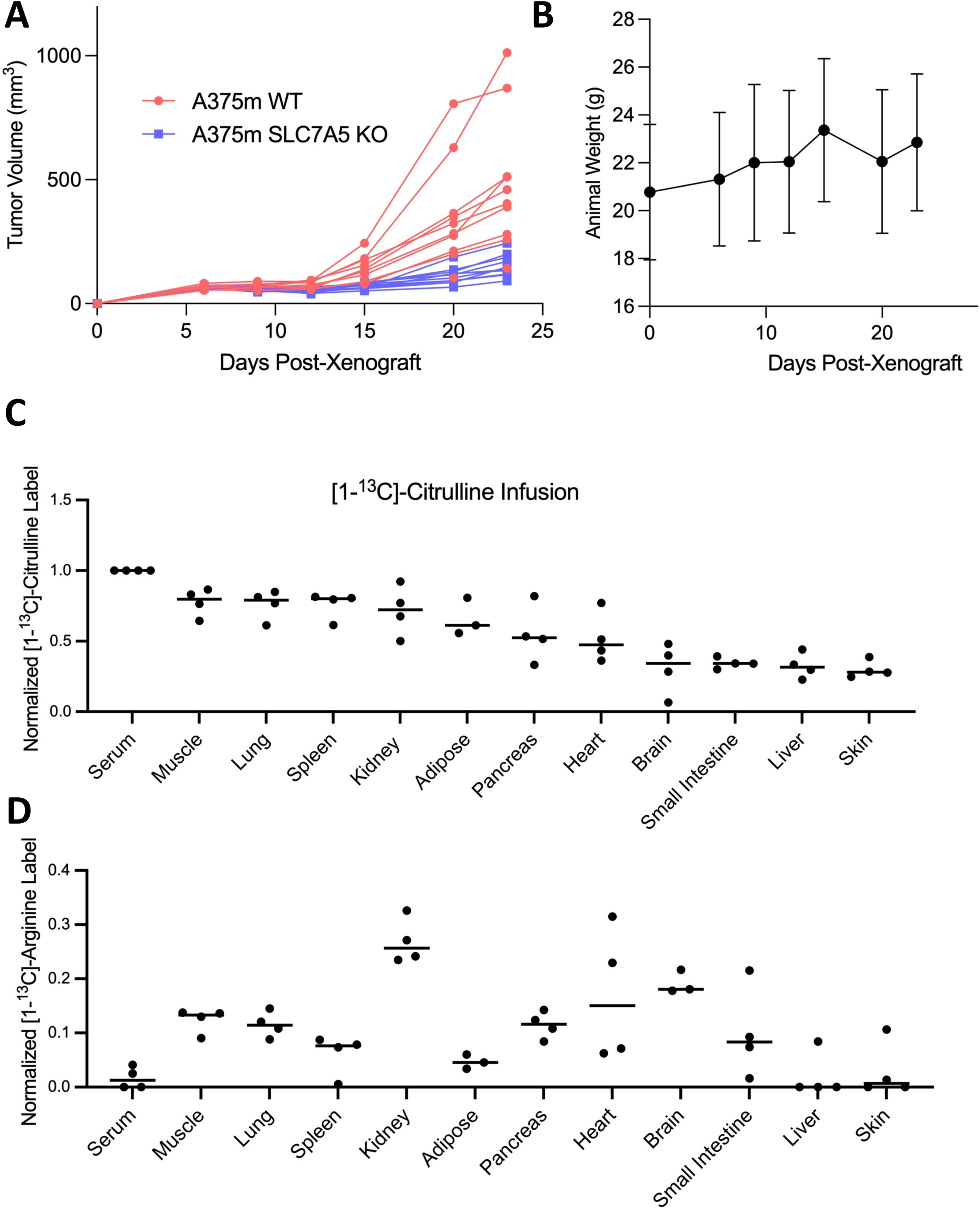
A: Tumor growth data of individual tumors as shown in Fig. 6B. n = 10 per color. B: Mouse weight in grams, corresponding to mice in Fig. 6B and Fig. S6A. C: Serum and tissue isotope labeling of citrulline after the 3-hour [1-^13^C]-Cit infusion. Data is normalized to the serum citrulline isotope label in each mouse. (mean ± SD, n = 4). D: Serum and tissue isotope labeling of arginine from citrulline after the 3-hour [1-^13^C]-Cit infusion. Data is expressed as a fraction of arginine label over citrulline label. (mean ± SD, n = 4).

## KEY RESOURCES TABLE

| REAGENT or RESOURCE | SOURCE | IDENTIFIER |
| --- | --- | --- |
| **Antibodies** |  |  |
| Rabbit polyclonal anti-LAT1(SLC7A5) | Cell Signaling Technology | Cat#5347; RRID: AB_10695104 |
| Rabbit polyclonal anti-PARP | Cell Signaling Technology | Cat#9542; RRID: AB_2160739 |
| Rabbit monoclonal anti-Cleaved PARP | Cell Signaling Technology | Cat#5625; RRID: AB_10699459 |
| Rabbit monoclonal anti-Caspase 3 | Cell Signaling Technology | Cat#14220; RRID: AB_2798429 |
| Rabbit monoclonal anti-Cleaved Caspase 3 | Cell Signaling Technology | Cat#9664; RRID: AB_2070042 |
| Rabbit monoclonal anti-ATF4 | Cell Signaling Technology | Cat#11815; RRID: AB_2616025 |
| Mouse monoclonal β-Actin-Peroxidase | Sigma Aldrich | Cat#A3854; RRID: AB_262011 |
| Goat Anti-Rabbit IgG-HRP | ThermoFisher | Cat#32260; RRID: AB_1965959 |
| **Bacterial and virus strains** |  |  |
| Invitrogen™ One Shot™ Stbl3™ Chemically Competent *E. coli* | ThermoFisher | Cat#C737303 |
| **Chemicals, peptides, and recombinant proteins** |  |  |
| RPMI | ThermoFisher | Cat#11875 |
| DMEM | ThermoFisher | Cat#11965 |
| SOC Media | New England Biolabs | Cat#B9020S |
| Fetal Bovine Serum | ThermoFisher | Cat#10437028 |
| Dialyzed Fetal Bovine Serum | Sigma-Aldrich | Cat#F0392 |
| Packed Cell Volume (PCV) Tubes | MidSci | Cat#TP87005 |
| Trypsin-EDTA w/ phenol red | ThermoFisher | Cat#25200056 |
| L-Arginine monohydrochloride | Sigma-Aldrich | Cat#A5131 |
| L-Citrulline | Sigma-Aldrich | Cat#C7629 |
| L-Phenylalanine | Sigma-Aldrich | Cat#P5482 |
| L-Leucine | Sigma-Aldrich | Cat#L8000 |
| L-Lysine monohydrochloride | Sigma-Aldrich | Cat#L5626 |
| RPMI 1640 Media w/o L-Arginine, L-Leucine, L-Lysine (Powder) | US Biologicals | Cat#R8999-03A |
| RPMI 1640 Media w/o L-Glutamine, w/o Amino Acids, Glucose, Phenol Red (Powder) | US Biologicals | Cat#R9010-02 |
| GCN2iB | MedChem Express | Cat#HY-112654 |
| JPH203 | Selleck Chemical | Cat#S8667 |
| ABT-263 | AdooQ Bioscience | Cat#A10022 |
| Puromycin Dihydrochloride | ThermoFisher | Cat#A1113803 |
| Blasticidin S HCl, powder | ThermoFisher | Cat#R21001 |
| HPLC Grade Methanol | Fisher Scientific | Cat#A452-4 |
| HPLC Grade Water | Fisher Scientific | Cat#W5SK-4 |
| HPLC Grade Acetonitrile | Fisher Scientific | Cat#14 650 359 |
| RIPA Buffer | Cell Signaling | Cat#9806S |
| Phosphatase Inhibitor Cocktail Powder | Sigma-Aldrich | Cat#P2714-1BTL |
| Sodium Orthovanadate | Sigma-Aldrich | Cat#S6508 |
| Tween-20 | Sigma-Aldrich | Cat#P9416 |
| 4x Lammeli Sample Buffer | BioRad | Cat#1610747 |
| 2-Mercaptoethanol | Sigma-Aldrich | Cat#M6250 |
| Proteinase K | New England Biolabs | Cat#P8107S |
| Trypan Blue 0.4% | ThermoFisher | Cat#15250061 |
| PMSF | Cell Signaling | Cat#8553S |
| Polybrene | Sigma Aldrich | Cat#TR-1003 |
| Poly-D-Lysine | ThermoFisher | Cat#A3890401 |
| Lipofectamine 3000 Transfection Reagent | ThermoFisher | Cat#L3000008 |
| L-Amino Acid Diet | Research Diets | Cat#A10021Bi |
| L-Amino Acid Diet -Arginine | Research Diets | Cat#A10036i |
| qScript cDNA SuperMix | VWR | Cat#95048 |
| Resazurin sodium salt | Sigma-Aldrich | Cat#R7017 |
| SYBR Green Master Mix | ThermoFisher | Cat#A46109 |
| NEBNext Ultra II Q5 Master Mix | New England Biolabs | Cat#M0544L |
| [1-<sup>13</sup>C]Citulline | Cambridge Isotopes | Cat#CLM-4899 |
| [U-<sup>13</sup>C]Arginine | Cambridge Isotopes | Cat#CLM-2265 |
| XTremeGene 9 | Sigma-Aldrich | Cat#6365779001 |
| Phosphate Buffered Saline, pH 7.4 | ThermoFisher | Cat#10010031 |
| Critical commercial assays |  |  |
| Pierce BCA Protein Assay Kit | Thermo Scientific | Cat#23227 |
| Pierce Rapid Gold BCA Protein Assay Kit | Thermo Scientific | Cat#A52335 |
| Trans Blot Turbo RTA Transfer Kit | BioRad | Cat#1704270 |
| QIAprep Spin Miniprep Kit | Qiagen | Cat#27106 |
| RNAeasy Mini Kit | Qiagen | Cat#74104 |
| DNeasy Blood & Tissue Kit | Qiagen | Cat#69504 |
| QIAmp DNA Blood Maxi Kit | Qiagen | Cat#51192 |
| Mycoplasma PCR Detection Kit | ABM | Cat#G238 |
| BioRad Gels 4-20% | BioRad | Cat#4561096 |
| Experimental models: Cell lines |  |  |
| Human: A375m | Roh-Johnson Lab,<br>University of Utah | N/A |
| Human: HCT116 | Rabinowitz Lab, Princeton | N/A |
| Human: HCT15 | Rutter Lab, U of U | N/A |
| Human: HEK-293FT | Roh-Johnson Lab, U of U | N/A |
| Human: LN229 | Rabinowitz Lab, Princeton | N/A |
| Human: MCF7 | Rutter Lab, U of U | N/A |
| Human: MDA-MB-468 | Roh-Johnson Lab, U of U | N/A |
| Human: NCI-H460 | McMahon Lab, U of U | N/A |
| Mouse: EO771 | Hilgendorf Lab, U of U | N/A |
| Experimental models: Organisms/strains |  |  |
| NOD.Cg-*Rag1*<sup>tm1Mom</sup> *Il2rg*<sup>tm1Wjl</sup>/SzJ (NRG) | University of Utah<br>Preclinical Research<br>Resource Core | N/A |
| C57BL6 | Charles River | Cat#027 |
| Oligonucleotides |  |  |
| sgRNA for CRISPR KO and qPCR Oligonucleotides | This study | Table S1 |
| Recombinant DNA |  |  |
| Human ASS1 in pLX304 | DNASU | Cat#HSCD00438196 |
| Human SLC7A5 in pDONR221 | DNASU | Cat#HSCD00042452 |
| pLX307 Empty Vector | Addgene | Cat#41392 |
| Human SLC7A5 in pLX307 | This study | N/A |
| pSpCas9(BB)-2A-GFP (PX458)-ASS1_exon11 | This study | N/A |
| pSpCas9(BB)-2A-Puro (PX459) V2.0-SLC7A5_exon2 | This study | N/A |
| pSpCas9(BB)-2A-Puro (PX459) V2.0 | Addgene | Cat#62988 |
| pSpCas9(BB)-2A-GFP (PX458) | Addgene | Cat#48138 |
| Human CRISPR Metabolic Gene Knockout Library | Addgene | Cat#110066 |
| psPAX2 | Addgene | Cat#12260 |
| PMD2.G | Addgene | Cat#12259 |
| Software and algorithms |  |  |
| MAGeCK | GitHub | https://github.com/liu-lab-dfci/MAGeCK |
| GraphPad Prism 10 | GraphPad Software | https://www.graphpad.com/ |
| R Version 4.4.0 | R | https://cran.r-project.org/ |
| EL-MAVEN Software | Elucidata | PMID: 31119671 |
| AccuCor | GitHub | https://github.com/XiaoyangSu/AccuCor |
| Biorender | Biorender | https://www.biorender.com/ |
| Other |  |  |
| Countess 3 Automated Cell Counter | Invitrogen | Cat#AMQAX2000 |
| BioTek Synergy Neo2 Hybrid Multimode Reader | Agilent | N/A |
| BioRad ChemiDoc MP Imaging System | BioRad | Cat#12003154 |
| Cryomill | Retsch | N/A |
| LC480 PCR Lightcycler | Roche | Cat#05015278001 |

## Notes

https://github.com/kndunlap/SLC7A5-Manuscript

### Competing Interest Statement

The authors have declared no competing interest.

